# Orai1α and Orai1β more potently promote NFAT activation and transcription than Orai2 and Orai3

**DOI:** 10.64898/2026.07.10.737753

**Authors:** Ahmed Emam Abdelnaby, Yin-Hu Wang, J. Cory Benson, Isagani Perez, Boheng Shen, Maxwell McDermott, Miki Jishage, Amal T. Elhaw, Silvia Cruz-Rangel, Raphael Courjaret, Abdelaziz Belkadi, Fang Yu, Douglas S. Prado, Shuai Yuan, Ping Xin, Adam C. Straub, Francisco J. Schopfer, Nadine Hempel, William F. Hawse, David B. Beck, Khaled Machaca, Stefan Feske, Mohamed Trebak

**Affiliations:** University of Pittsburgh, Department of Pharmacology and Chemical Biology, Pittsburgh, PA, USA; Department of Pathology, New York University Grossman School of Medicine, New York, NY, USA; Department of Medicine, New York University Grossman School of Medicine, New York, NY, USA; UPMC Hillman Cancer Center, University of Pittsburgh School of Medicine, Pittsburgh, PA, USA; Department of Medicine, Division of Malignant Hematology and Medical Oncology, University of Pittsburgh School of Medicine, Pittsburgh, PA, USA; Calcium Signaling Group, Research Department, Weill Cornell Medicine Qatar, Education City, Qatar Foundation, Doha 24144, Qatar; and Department of Systems and Computational Biomedicine, Weill Cornell Medicine, New York, USA; Department of Immunology, University of Pittsburgh, Pittsburgh, PA, USA; Center for Systems Immunology, University of Pittsburgh, Pittsburgh, PA, USA

**Keywords:** CRAC channels, Orai isoforms, calcium signaling, NFAT, immune function, channelopathy

## Abstract

The Ca²⁺ release-activated Ca²⁺ (CRAC) channel mediates store-operated calcium entry (SOCE), a ubiquitous pathway essential for many cell types, including immune cells. Three Orai (Orai1, Orai2, Orai3) proteins constitute the plasma membrane pore-forming units of CRAC channels that are activated by the endoplasmic reticulum (ER) Ca^2+^-sensing STIM1 and STIM2 proteins when ER Ca^2+^ stores are depleted. Orai1/2/3 are differentially expressed across primary cells with discernible differences in their structures and biophysical properties. Further, Orai1 has two alternatively translated isoforms: the long mammalian-specific Orai1α and the 63-residue shorter Orai1β, which is evolutionarily older and conserved across vertebrates. Whether Orai1α/1β/2/3 at native expression levels produce unique cytosolic Ca²⁺ signatures that bias transcriptional responses through effectors like NFAT is unclear. Here we used HEK293 cells engineered to express one native Orai isoform and show that all Orai isoforms couple to NFAT1 and NFAT4 induction. The magnitude of NFAT1/4 induction tracked the magnitude of SOCE, with the following profile: Orai1β>Orai1α>>Orai2>Orai3. Near-native ectopic expression of either Orai1α or Orai1β in primary murine *Orai1^−/−^*CD4⁺ T cells restored SOCE, NFAT activation, cytokine production and promoted near identical transcriptional responses enriched for immune activation pathways. An analysis of genetic and clinical data of human individuals showed that homozygous null mutations selectively abolishing Orai1α are not associated with disease resembling CRAC channelopathy. Primary T cells from individuals homozygous or heterozygous for an Orai1α null mutation showed enhanced, rather than impaired, SOCE and NFAT induction. Collectively, our data indicate that Orai isoforms regulate NFAT activation and downstream transcription primarily through the magnitude of SOCE they support, with Orai1α and Orai1β being the most potent isoforms coupling SOCE to NFAT responses.

## Introduction

Calcium (Ca²⁺) is a ubiquitous second messenger with central roles in signal transduction (1). One of the predominant Ca²⁺ influx pathways into the cytosol of non-excitable cells is store-operated Ca²⁺ entry (SOCE) (2–6). SOCE is mediated by Ca²⁺ release–activated Ca²⁺ (CRAC) channels (7, 8), which are formed by the pore-forming subunits Orai1/2/3 (9), and activated by the endoplasmic reticulum (ER)-resident Ca²⁺ sensors STIM1 and STIM2 (10, 11) when ER Ca^2+^ stores are depleted. Each of the three Orai and the two STIM proteins is encoded by a distinct gene. Further, Orai1 exists as two alternatively translated variants, a longer, mammalian-specific isoform (Orai1α) and shorter isoform (Orai1β) lacking the first N-terminal 63 amino acids (12, 13). Upon ER Ca²⁺ store depletion, STIM proteins oligomerize, move to plasma membrane-ER contact sites and gate Orai channels, enabling Ca²⁺ influx from the extracellular space to replenish ER stores and to activate cellular signaling pathways that couple to transcriptional and metabolic activities (14). The Ca^2+^ microdomains generated at the cytosolic mouth of CRAC channels are decoded by a network of Ca²⁺-responsive signaling pathways, culminating in the activation of multiple transcription factors (15, 16). Among these, nuclear factor of activated T cells (NFAT) is well-characterized as a canonical downstream target for SOCE (17–19). In resting cells, NFAT is retained in the cytoplasm through phosphorylation that masks its nuclear localization sequence. Ca²⁺ influx through Orai channels activates the phosphatase calcineurin, which dephosphorylates NFAT, enabling its nuclear translocation and thereby linking CRAC-mediated Ca²⁺ influx to gene programs that orchestrate diverse biological processes, including proliferation, immune activation, cytokine production, and metabolic regulation (14, 20–23). The NFAT family comprises five members, four of which (NFAT1–4) are activated in a Ca²⁺-dependent manner. NFAT1 (NFATc2) and NFAT4 (NFATc3), the two family members examined here, have both shared and context-dependent physiological functions. In T cells, NFAT1 contributes to peripheral T-cell activation, cytokine production, differentiation, and immune homeostasis, whereas NFAT4 has prominent roles in thymocyte development and positive selection; beyond the immune system, both factors participate in tissue-specific programs, including skeletal muscle and cardiovascular biology (22, 24, 25). NFAT1 and NFAT4 also display distinct activation kinetics and Ca²⁺ sensitivities: NFAT4 nuclear translocation occurs rapidly in response to modest cytosolic Ca²⁺ rises and is exported quickly from the nucleus, whereas NFAT1 requires stronger, more prolonged cytosolic Ca²⁺ signals for activation and has slower nuclear export kinetics (26–28).

Orai homologues share a common activation mechanism, but they diverge in tissue expression, regulation and biophysical properties, including their susceptibility to fast Ca²⁺-dependent inactivation (CDI), from the high fast CDI of Orai3 to the minimal fast CDI of Orai1β (12, 29, 30). These differences, together with their ability to assemble in varying stoichiometries, potentially enable native CRAC channels to fine-tune Ca²⁺ signal amplitude, duration and dynamics, thereby broadening the bandwidth of Ca²⁺ signaling events (20, 31–33). For instance, knockdown of Orai2 or Orai3 enhances SOCE, suggesting that these isoforms act as regulatory brakes on CRAC channel activity (32, 34, 35). Yet, how the unique Ca²⁺ signatures of individual Orai isoforms shape the transcriptional landscape remains largely unknown. Further, whether Orai1α and Orai1β isoforms are each capable of supporting SOCE-dependent transcriptional and functional responses in a physiological context such as in primary CD4⁺ T cells has not been examined. The longer Orai1α isoform contains several unique motifs and residues shown to affect SOCE itself and SOCE-dependent downstream signaling. Substitution of serines 27 and 30, putative protein kinase C phosphorylation sites unique to Orai1α, resulted in increased SOCE suggesting that Orai1α phosphorylation on S27/S30 regulates CRAC channel function (36). Moreover, phosphorylation of serine 34 following Ca^2+^ influx through Orai1α was shown to mediate CRAC channel inactivation in a cyclic AMP and protein kinase A-dependent manner (37). Orai1α contains a caveolin consensus site between residues 52-60 that have been shown to mediates its internalization from the plasma membrane during oocyte meiosis (38). Previous studies also proposed that residues 39–59 of Orai1α can bind the scaffolding protein AKAP79, thus coupling SOCE to calcineurin and NFAT activation (39, 40).

Here we set out to analyze all four Orai isoforms (Orai1α/1β/2/3) at their near native levels of expression in HEK293 and primary CD4^+^ T cells for their ability to mediate SOCE, activate NFAT isoforms, and regulate gene expression. Using HEK293 cells natively expressing only one Orai homologue, we found that Orai1 is the principal driver of NFAT1/4 activation whereas Orai2 and Orai3 less efficiently supported NFAT1/4 nuclear translocation. Comparing Orai1α and Orai1β isoforms, we found that both isoforms can promote NFAT nuclear translocation in HEK293 cells across diverse activation conditions with Orai1β mediating significantly bigger NFAT nuclear translocation compared to Orai1α. Expression of Orai1α and Orai1β at near native levels in primary *Orai1^−/−^* CD4⁺ T cells showed that both restored SOCE, NFAT translocation, cytokine production and gene expression without appreciable differences. An analysis of genetic and clinical data showed that human individuals with homozygous null mutations unique to Orai1α who only express Orai1β protein do not suffer from CRAC channelopathy syndrome in contrast to patients with null mutations in residues shared by both Orai1α and Orai1β (41). Moreover, the analysis of T cells from patients heterozygous or homozygous for an Orai1α null mutation showed increased SOCE and NFAT1 activation compared to healthy controls expressing both isoforms. Collectively, our findings demonstrate that all three Orai homologues and both Orai1 isoforms can mediate NFAT activation and cell functions, albeit with different efficiencies that correlate with the magnitude of SOCE through these channels.

## Results

### Native Orai1 promotes more efficient NFAT1/4 nuclear translocation upon maximal store depletion compared to native Orai2 and Orai3

Ca²⁺ influx through Orai channels generates spatially restricted cytosolic Ca^2+^ microdomains that couple SOCE to transcriptional regulation, controlling multiple gene expression programs (15, 22, 42–45). To dissect Orai homologue-specific contributions to NFAT activation, and ultimately gene expression, we used CRISPR/Cas9-engineered HEK293 cell clones lacking two Orai isoforms (double knockout, Orai-DKO), thereby retaining expression of the third Orai isoform at endogenous expression levels. These knockout models were previously validated at the genomic and transcript levels, with no compensatory changes in expression of STIM1, STIM2 or IP_3_ receptors (32, 37). NFAT1 and NFAT4 were shown previously to require different types of Ca²⁺ signals in terms of magnitude and source of Ca²⁺ (32, 37, 46). We therefore expressed NFAT1-GFP and NFAT4-GFP reporters in each DKO cell line and tracked their localization by live imaging following stimulation with 2 μM thapsigargin in the presence of extracellular Ca²⁺ to induce strong, sustained SOCE. Nuclear NFAT accumulation was quantified as the nuclear-to-cytoplasmic GFP ratio over time on a cell-by-cell basis. Native Orai1 induced the most robust NFAT nuclear translocation compared to Orai2 or Orai3 in both the maximum level and kinetics of translocation (**Fig. 1A-H**). Orai1 was more efficient than Orai2 and Orai3 in inducing the translocation of both NFAT1 and NFAT4. Of note, Orai2 was more efficient at inducing NFAT4 nuclear translocation compared to Orai3 in cells stimulated with thapsigargin. These findings demonstrate that under conditions of maximal SOCE induction, Orai1 is the main driver of both NFAT1 and NFAT4 nuclear translocation.

**Figure 1.**
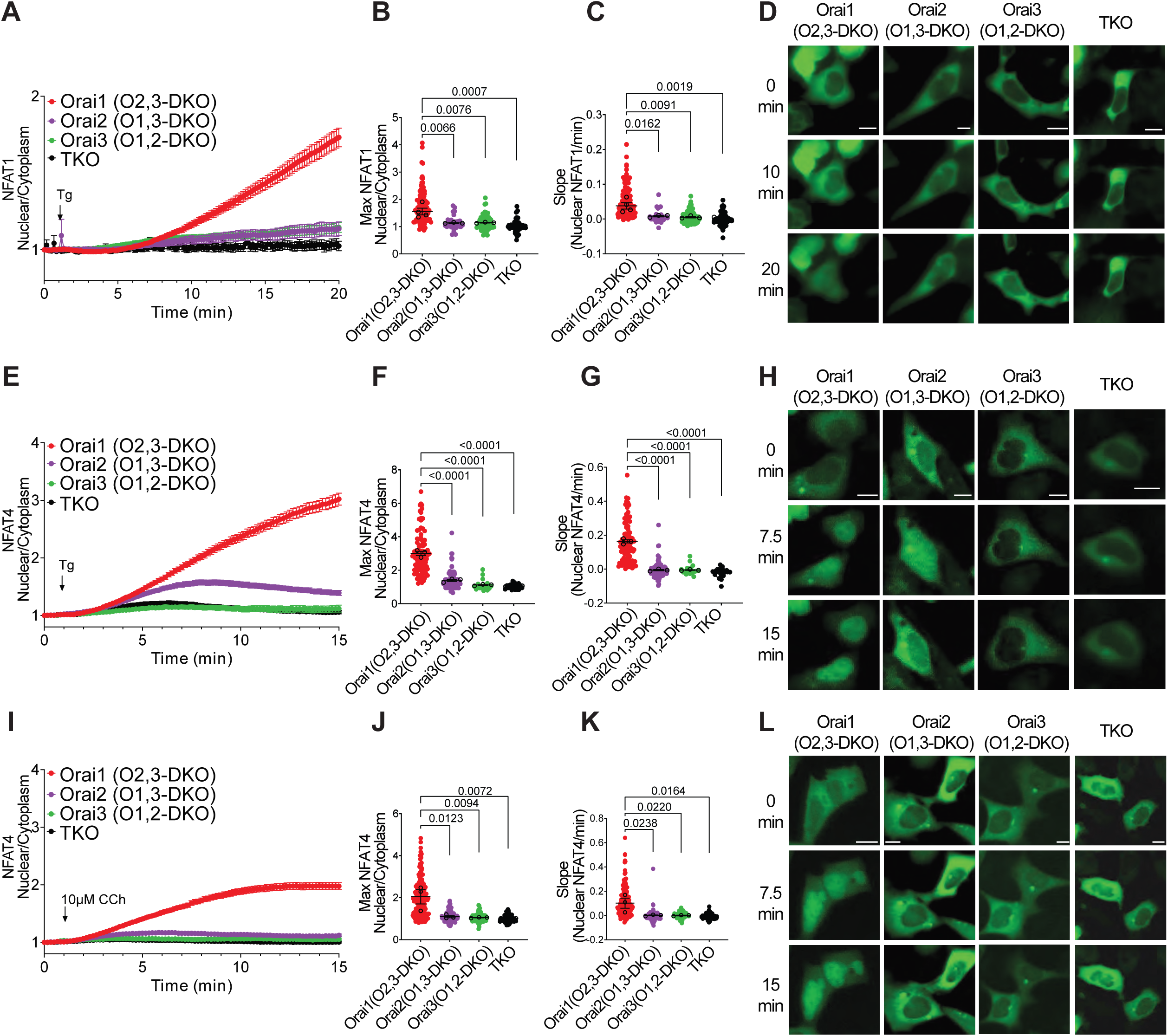
Native Orai1 is the primary driver of NFAT1/4 nuclear localization. NFAT1-GFP **(A–D)** and NFAT4-GFP **(E-L)** nuclear translocation in HEK293 Orai double knockout (DKO) cells stimulated with either 2 μM thapsigargin (Tg; **A-H**) or 10 μM CCh (**I-L**) in the presence of 2 mM extracellular Ca²⁺. Panels represent time-lapse traces of NFAT nuclear/cytoplasmic fluorescence ratios (**A, E, I)**, quantification of maximal NFAT nuclear/cytoplasmic fluorescence ratios (**B, F, J**), quantification of NFAT translocation rates (**C, G, K**) and representative images of NFAT-GFP nuclear localization at the indicated times post stimulation (**D, H, L**). For quantification of NFAT translocation rates, slopes were calculated from 8–15 min (**C**), 5–15 min (**G**), and 5–10 min (**K**). Data are expressed as mean ± SEM of the nuclear/cytoplasmic GFP fluorescence ratio. Individual-cell measurements are shown together with the mean of each biological replicate. Statistical analyses were performed using the biological-replicate means by ordinary one-way ANOVA followed by Dunnett’s multiple-comparisons test. Adjusted P values for the indicated comparisons are shown. A direct comparison of maximal NFAT4 nuclear translocation between Orai2 and Orai3 following thapsigargin stimulation was performed using Welch’s t test (P = 0.0358) (F). Scale bar, 10 μm.

### Orai1 supports robust NFAT4 nuclear localization upon physiological receptor stimulation

To determine the differential roles of Orai homologues in NFAT nuclear translocation under more physiological conditions of activation, we next stimulated cells with the G-protein coupled receptor (GPCR) agonist carbachol (CCh) at 10 µM, a relatively low dose that typically generates robust oscillations in HEK293 cells (31, 32, 47). Although these oscillations are initiated by IP₃R-mediated release, CRAC channel activity is essential for sustaining them and for driving downstream transcriptional programs (31, 32). Because NFAT1 nuclear translocation requires robust and sustained CRAC channel activity, low concentrations of GPCR agonists do not cause detectable activation of NFAT1. Consistent with this, we previously showed that 10 µM CCh fails to induce NFAT1 nuclear translocation in HEK293 cells (37). Therefore, we focused on NFAT4. Using the homologue-specific Orai-DKO cell lines, we expressed NFAT4-GFP and monitored its localization by live imaging following stimulation with 10 µM CCh. Native Orai1 mediated robust NFAT4 nuclear accumulation compared to Orai2 and Orai3 (**Fig. 1I–L**). These findings demonstrate that Orai1 is the principal Orai family member coupling SOCE to NFAT4 nuclear localization after receptor-mediated stimulation of SOCE.

### Orai1α and Orai1β both promote NFAT1/4 nuclear localization

We next asked whether the two alternatively translated Orai1 isoforms, Orai1α and Orai1β, differ in their ability to activate NFAT1/4 nuclear translocation. Orai1β arises from an alternative translation start site (TSS) at methionine 64, producing a shorter isoform lacking the distal N-terminal 63 amino acids (**Fig. 2A**) (12, 13). To directly compare the effects of both Orai1 isoforms, we used HEK293 triple Orai knockout (Orai-TKO) cells lacking all three Orai homologues and reintroduced either Orai1α or Orai1β cDNA plasmid. To achieve near-native levels of Orai1α and Orai1β, their expression was placed under the control of the relatively “weak” chicken thymidine kinase (TK) promoter. This system offers several advantages over conventional plasmid overexpression driven by the strong cytomegalovirus (CMV) promoter. Using both immunofluorescence and Western blotting, we show by comparison to CMV-driven plasmids, TK-driven plasmids achieve near-endogenous levels of Orai isoform expression (**Fig. S1**). These TK-driven near-physiological expression levels better preserve Orai-STIM stoichiometry, thereby eliminating the need for STIM1 co-expression and avoiding artifacts associated with excessive cDNA loading (37). This is important because overexpression of Orai/STIM genes has been shown to obscure Orai isoform-specific differences and alter the effects of endogenous SOCE regulators (12, 48, 49).

**Figure 2.**
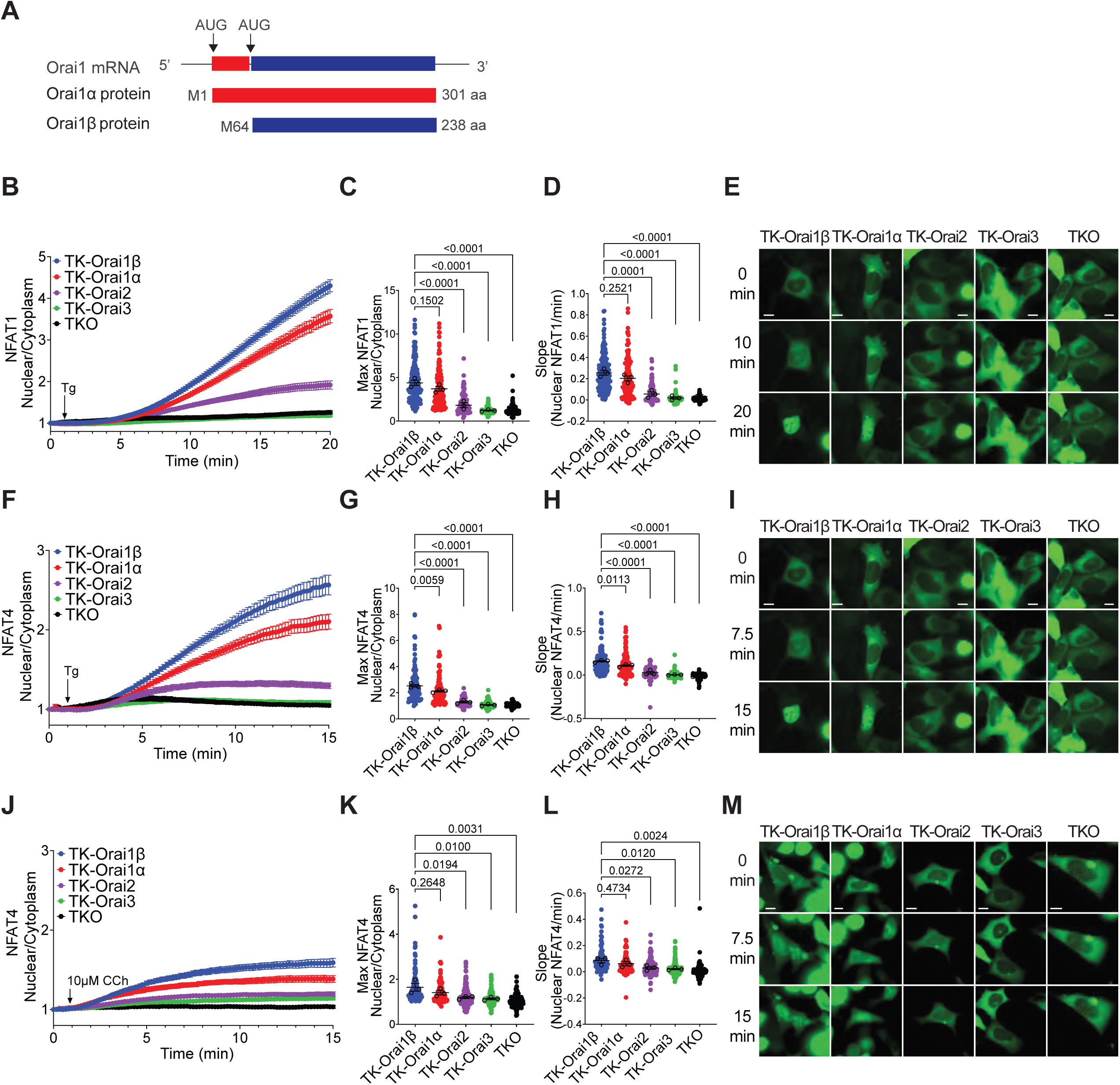
Both Orai1α and Orai1β drive NFAT1/4 nuclear localization. **(A)** Schematic illustrating alternative translation initiation of Orai1 mRNA, producing Orai1α (301 aa) and Orai1β (238 aa). NFAT1-GFP (**B-E**) and NFAT4-GFP (**F-M**) nuclear translocation in HEK293 Orai triple knockout (TKO) cells reconstituted with native levels of Orai1α and Orai1β isoforms expressed under the control of the chicken thymidine kinase (TK) promoter. Cells were stimulated with either 2 μM thapsigargin (Tg, **B-I**) or 10 μM CCh (**J-M**) in the presence of 2 mM extracellular Ca²⁺. Panels represent time-lapse traces of NFAT nuclear/cytoplasmic fluorescence ratios (**B, F, J**), quantification of maximal NFAT nuclear/cytoplasmic fluorescence ratios (**C, G, K**), quantification of NFAT translocation rates (**D, H, L**) and representative images of NFAT-GFP nuclear localization at the indicated times post stimulation (**E, I, M**). For the quantification of NFAT translocation rates, slopes were calculated from 10–15 min (**D**), 5–10 min (**H**), and 2–5 min (**L**). Data are expressed as mean ± SEM of the nuclear/cytoplasmic GFP fluorescence ratio. Individual-cell measurements are shown together with the mean of each biological replicate. Statistical analyses were performed using the biological-replicate means by ordinary one-way ANOVA followed by Dunnett’s multiple-comparisons test. Adjusted P values for the indicated comparisons are shown. Scale bar, 10 μm.

With comparable expression levels established for Orai1α, Orai1β, Orai2 and Orai3 (**Fig. S1D**), we compared the ability of these four channel proteins to promote the activation of NFAT1 and NFAT4 in response to weak and strong induction of SOCE. NFAT1-GFP and NFAT4-GFP reporter assays revealed that both Orai1α and Orai1β efficiently supported NFAT1 and NFAT4 nuclear translocation upon stimulation with 2 μM thapsigargin (**Fig. 2B–I**). A quantitative analysis confirmed significant increases in both the maximal nuclear/cytoplasmic ratio and the slope of translocation kinetics for both Orai1α and Orai1β isoforms compared to Orai2, Orai3, or Orai-TKO controls. Of note, Orai1β induced significantly greater and faster NFAT4 nuclear translocation than Orai1α following thapsigargin stimulation (**Fig. 2G, H**), which is consistent with the enhanced fast Ca^2+^-dependent inactivation (CDI) of Orai1α compared to Orai1β (12). Using more physiological activation conditions by stimulation with 10 µM CCh yielded similar results. Orai1α and Orai1β both effectively promoted NFAT4 nuclear localization (**Fig. 2J–M**). These results demonstrate that both Orai1α and Orai1β effectively couple SOCE to NFAT1/4 activation under conditions of maximal store depletion and physiological receptor stimulation.

### The magnitude of NFAT nuclear translocation is commensurate with SOCE levels and is not Orai isoform specific

To determine whether the magnitude and kinetics of NFAT activation reflect differences in Ca²⁺ entry mediated by Orai1α, Orai1β, Orai2, and Orai3, we directly measured SOCE in Orai-TKO cells ectopically expressing individual Orai homologues and isoforms using TK promoter-driven constructs. Equivalent expression levels of YFP-tagged proteins in individual cells were validated (**Fig. S1B-D**). SOCE, which in this situation is supported by native STIM proteins, was measured following maximal store depletion with thapsigargin. These recordings showed that Orai1β expression resulted in the largest SOCE magnitude, followed by Orai1α, Orai2, and Orai3 (**Fig. 3A–B**). This rank order tracked closely with NFAT nuclear translocation (**Fig. 2**), suggesting that NFAT responses reflect the relative Ca²⁺ influx mediated by each Orai isoform. Consistent with this conclusion, mean maximal NFAT1 and NFAT4 responses across the five Orai conditions showed a positive correlation with mean maximal SOCE (R² = 0.93 and 0.94, respectively) (**Fig. S2**).

**Figure 3.**
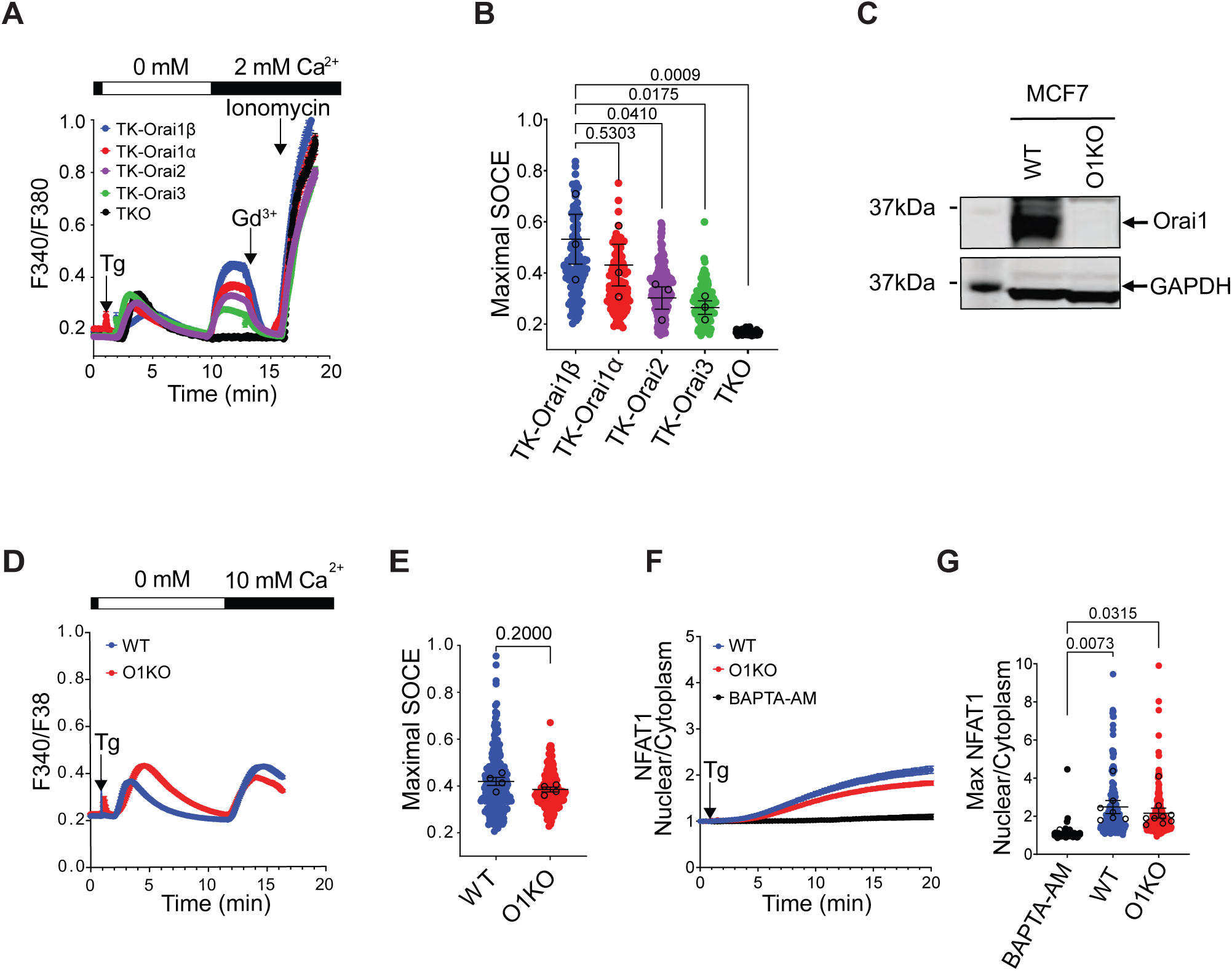
The magnitude of NFAT activation mirrors that of Ca²⁺ influx driven by each Orai isoform. **(A)** SOCE traces in HEK293 Orai-TKO cells reconstituted with each of the four Orai isoforms expressed from TK-driven constructs at near endogenous levels and C-terminally tagged with YFP. Recordings were performed on cells with equivalent levels of YFP fluorescence. Cells were stimulated with 2 μM thapsigargin (Tg) in Ca²⁺-free medium, followed by re-addition of 2 mM extracellular Ca²⁺; 5 μM Gd³⁺ was applied to block CRAC currents and 10 μM Ionomycin was added at the end of the recordings to induce SOCE-independent maximal Ca^2+^ influx. **(B)** Quantification of maximal SOCE amplitude mediated by each Orai isoform compared to mock-transfected HEK293 TKO cells. **(C)** Immunoblot confirming Orai1 knockout in MCF7 cells (MCF7-O1KO) compared with wildtype (WT) control MCF7 cells; GAPDH served as a loading control. **(D)** SOCE recordings in WT MCF7 cells and MCF7-O1KO cells. **(E)** Quantification of SOCE peak in WT MCF7 cells and MCF7-O1KO cells from D. **(F)** Time lapse of NFAT1 nuclear translocation following thapsigargin (Tg) stimulation in MCF7 WT, MCF7-O1KO, and MCF7 WT cells treated with 20 μM BAPTA-AM. **(G)** Quantification of maximal NFAT1 nuclear/cytoplasmic ratios in each condition from F. Data are expressed as mean ± SEM of the nuclear/cytoplasmic GFP fluorescence ratio. Individual-cell measurements are shown together with the mean of each biological replicate. Statistical analyses were performed using the biological-replicate means by ordinary one-way ANOVA followed by Dunnett’s multiple-comparisons test. Adjusted P values for the indicated comparisons are shown. Scale bar, 10 μm.

Our data thus far demonstrate that Orai2 and Orai3 channels support NFAT1/4 translocation although it was rather modest. If NFAT activation is mainly governed by the magnitude of SOCE, we reasoned that the homologue mediating the lowest SOCE, Orai3, might still support robust NFAT activation if it is highly expressed under native conditions. Thus, we examined MCF7 breast cancer cells, in which Orai3 is highly expressed and is the predominant functional Orai channel mediating SOCE. In MCF7 breast cancer cells, Orai3-mediated SOCE contributes to breast cancer cell proliferation and survival (50–55). To ensure that the bulk of SOCE in MCF7 cells is mediated by Orai3 and to rule out contributions from Orai1 to SOCE, we deleted Orai1 using CRISPR/Cas9 gene editing (MCF7-O1KO) (29) (**Fig. 3C**). Consistent with our previous findings, SOCE was only slightly reduced in the absence of Orai1 (**Fig. 3D-E**), confirming that Orai3 is the major contributor to CRAC channel activity in MCF7 cells (50, 53, 54, 56). We then monitored NFAT1 nuclear translocation and found that following thapsigargin stimulation MCF7-O1KO cells sustained robust NFAT1 activation comparable to that of wildtype MCF7 cells (**Fig. 3F–G**). As a negative control, wildtype MCF7 cells pretreated with 20 μM BAPTA-AM to chelate cytosolic Ca^2+^ showed no NFAT1 nuclear translocation (**Fig. 3F-G**). These data demonstrate that abundant, native Orai3 can effectively promote NFAT1 nuclear translocation, supporting the conclusion that NFAT1 activation scales with the capacity of individual Orai homologues to mediate SOCE.

### Orai1α- and Orai1β-dependent NFAT activation is governed by Ca^2+^ microdomains at the mouth of the channel

Previous studies have shown that Ca²⁺ signals can be decoded by Ca²⁺-binding proteins in the vicinity of Ca²⁺ channels and relayed to the nucleus to achieve distinct transcriptional patterns (57, 58). Given the dominant role of Orai1α and Orai1β in NFAT activation in most cell types, we asked whether the SOCE-dependent nuclear translocation of NFAT1 is regulated by the magnitude of the global Ca^2+^ signal or localized Ca²⁺ signals generated at the mouth of Orai1 channels. To this end, we studied the impact of slow and fast Ca²⁺-binding chelators, namely membrane-permeant forms of EGTA-AM and BAPTA-AM respectively, on Orai1 isoform dependent SOCE and subsequent NFAT1 nuclear translocation. Both chelators have similar affinities for Ca²⁺ but differ in their binding rates. EGTA’s slower binding kinetics creates a region of relatively unbuffered intracellular Ca²⁺ concentration at the vicinity of individual Ca²⁺ channels. On the other hand, BAPTA’s faster binding rate results in a sharp decrease in Ca^2+^ concentrations within a small distance away from a Ca²⁺ channel (59, 60). Therefore, BAPTA is likely to inhibit both local and global Ca²⁺ signals, while EGTA suppresses only the global Ca²⁺ signal (**Fig. 4A**). These differential effects have been extensively employed to investigate local channel-effector coupling in diverse contexts (57, 58, 61–63). We pre-loaded Orai-TKO HEK293 cells expressing either Orai1α or Orai1β with EGTA-AM or BAPTA-AM and analyzed SOCE and NFAT1-GFP nuclear translocation. Both EGTA and BAPTA completely suppressed the global Ca²⁺ increase after thapsigargin stimulation of cells expressing either Orai1α or Orai1β (**Fig. 4B-C**). By contrast, only BAPTA, but not EGTA, strongly interfered with the nuclear translocation of NFAT1 in response to thapsigargin stimulation (**Fig. 4D-I**). These findings, which are consistent with earlier reports (28, 39, 64, 65), indicate that NFAT1 activation is regulated by localized Ca²⁺ signals in the vicinity of the CRAC channel pore. Of note, the inhibitory effect of BAPTA on NFAT1-GFP translocation was observed in cells expressing either Orai1α or Orai1β indicating that it is not isoform specific.

**Figure 4.**
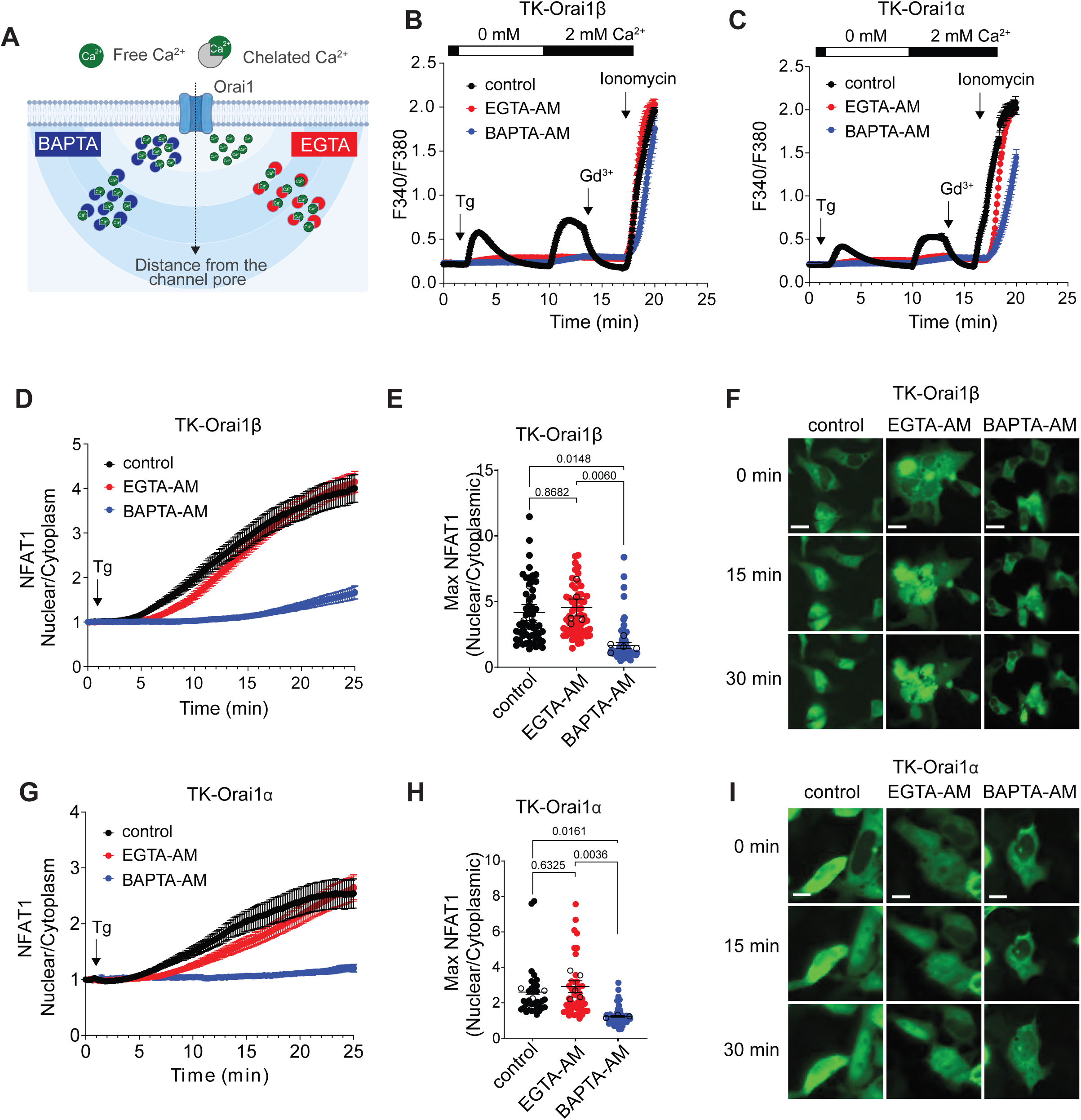
NFAT activation by Orai1α and Orai1β depends on localized Ca²⁺ signals near the channel pore. **(A)** Schematic illustrating the fast Ca²⁺ chelator BAPTA and the slower chelator EGTA differentially buffer local versus global Ca²⁺ signals. **(B–C)** SOCE recordings in HEK293 Orai-TKO cells expressing native levels of Orai1β **(B)** or Orai1α **(C)** driven by the TK promoter, showing that both chelators reduced global Ca²⁺ influx. **(D–I)** NFAT1 nuclear translocation in cells reconstituted with either Orai1β **(D-F)** or Orai1α **(G-I)**, which was suppressed by BAPTA-AM but not by EGTA-AM loading. **(D, G)** Time course of NFAT1 nuclear/cytoplasmic fluorescence ratio following thapsigargin (Tg) stimulation in both conditions. **(E, H)** Quantification of maximal NFAT1 nuclear translocation. **(F, I)** Representative images of NFAT1-GFP nuclear localization at indicated times after Tg stimulation in control, EGTA-AM–treated, and BAPTA-AM–treated cells. Data are expressed as mean ± SEM of the nuclear/cytoplasmic GFP fluorescence ratio. Individual-cell measurements are shown together with the mean of each biological replicate. Statistical analyses were performed using the biological-replicate means by ordinary one-way ANOVA followed by Tukey’s multiple-comparisons test. Adjusted P values for the indicated comparisons are shown. Scale bar, 10 μm.

The mechanisms underlying regulation of NFAT activation by local Ca^2+^ signals mediated by Orai channels are not well understood. In neurons, the binding of the Ca^2+^ sensor calmodulin to voltage-gated Ca^2+^ channels is a known mechanism to recruit calmodulin kinases and activate the transcription factor CREB near the channel pore (57, 66). A recent study showed that Orai1 may use a similar mechanism that includes binding of the adaptor AKAP79 to a 21 amino acid region (39–59) in the N-terminus of Orai1α, thus facilitating AKAP79-dependent recruitment of calcineurin to Orai1 channels and NFAT1 activation (40). Because the AKAP79-binding region is missing in Orai1β, we hypothesized that NFAT1 activation would be reduced in cells expressing Orai1β compared to Orai1α. Contrary to this hypothesis, nuclear translocation of NFAT1 and NFAT4 was comparable in Orai1β and Orai1α expressing cells (**Fig. 2**). Collectively, our findings indicate that while local Ca^2+^ signals near the CRAC channel pore mediate the activation of NFAT, they are unlikely to be decoded by the N-terminal region unique to Orai1α.

### Both Orai1α and Orai1β promote NFAT activation and cytokine production in primary CD4^+^ T cells

We next sought to determine if Orai1α and Orai1β show differences in their ability to promote NFAT nuclear translocation and NFAT-dependent gene expression in cells in which Orai1 is essential for function. Orai1 and NFAT are indispensable for T cell activation, cytokine production, and effector functions (16, 67). Native Orai1α and Orai1β proteins can be resolved using Western blotting of deglycosylated proteins samples as described previously (12, 13). Primary naïve CD4^+^ T cells as well as CD4^+^ T cells differentiated into either T helper (Th) 1, Th2, regulatory (Treg), non-pathogenic Th17 (npTh17) or pathogenic Th17 (pTh17) express both Orai1α and Orai1β isoforms with varying ratios (**Fig. S3**), making T cells an excellent model to probe the functional significance of Orai1α and Orai1β signaling. To this end, we isolated primary murine CD4^+^ T cells from the spleens of *Orai1^fl/fl^ Cd4Cre* mice (*Orai1^CD4^*) with T cell specific deletion of both Orai1α and Orai1β (68) and retrovirally transduced them with either Orai1α or Orai1β in plasmids containing the fluorescent reporter Ametrine (**Fig. S4A**). Ametrine (Amt) fluorescence correlated well with Orai1α and Orai1β expression levels detected by antibody staining (**Fig. S4B**). This allowed us to use Amt reporter fluorescence as a proxy for Orai1 levels, which is important to ensure similar expression levels of Orai1α and Orai1β. Because strong ectopic Orai1 expression suppresses SOCE – likely due to an increased Orai1:STIM1 ratio (69, 70) – we gated on Amt^high^ and Amt^low^ cells to control for Orai1 expression levels (**Fig. S4C-D**). Measurements of intracellular Ca^2+^ concentrations using Indo-1 showed that *Orai1^CD4^* T cells expressing high levels of either Orai1α or Orai1β (Amt^high^) had low thapsigargin-induced SOCE similar to cells expressing EV control (**Fig. S4F-G**), consistent with the previously reported suppressive effects of supraphysiological Orai1 expression on SOCE. By contrast, T cells expressing low levels of Orai1 (Amt^low^) showed increased SOCE compared to EV-transduced cells (**Fig. 5A**).

**Figure 5.**
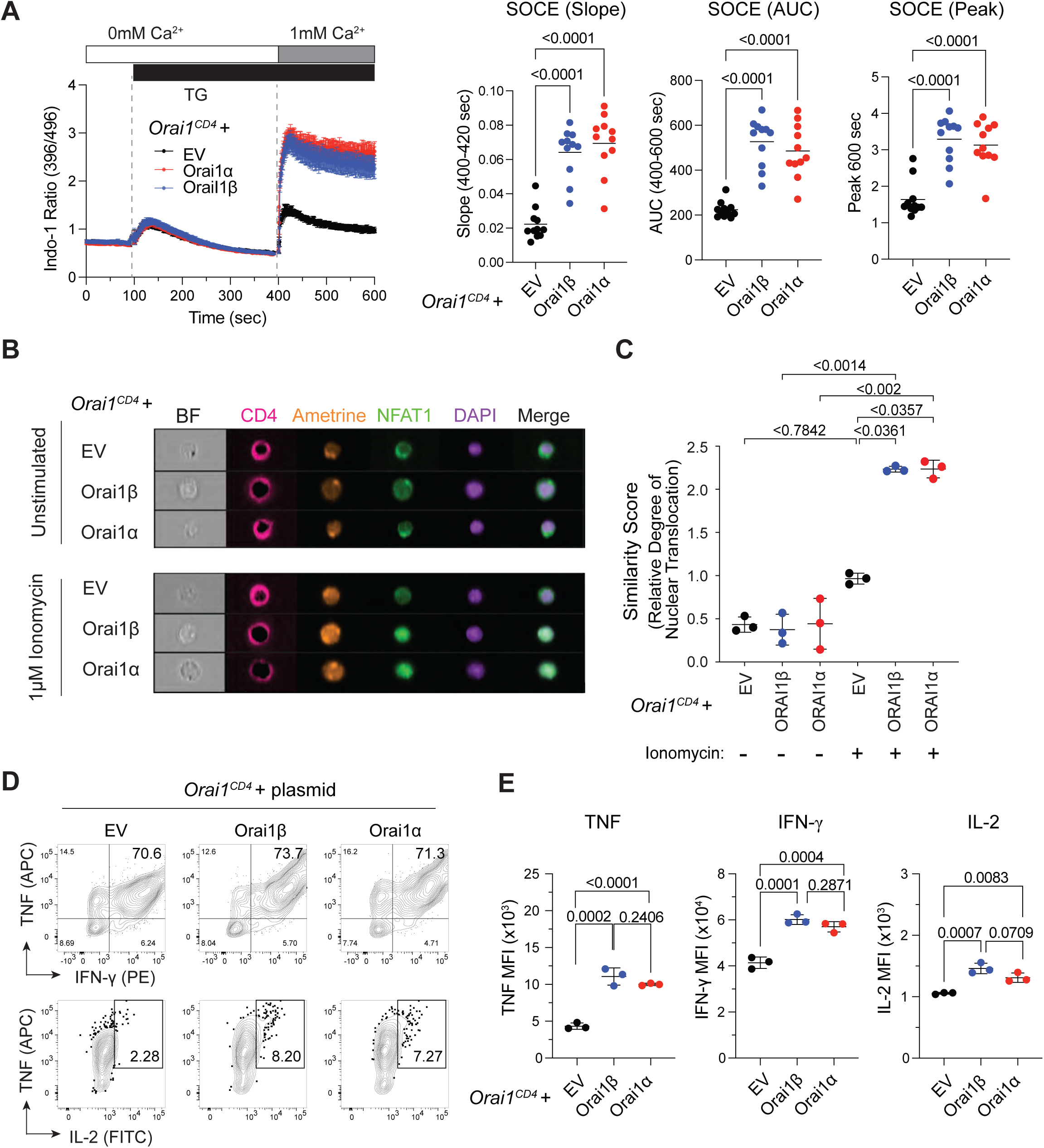
Both Orai1α and Orai1β support NFAT signaling and cytokine production in primary CD4⁺ T cells. CD4⁺ T cells from *Orai1^fl/fl^ Cd4^Cre^* mice were retrovirally transduced with bicistronic vectors encoding either Orai1α or Orai1β followed by an IRES-Ametrine reporter, or with an empty vector (EV) control. Cells were polarized into Th1 cells for an additional 2 days. (A) Analysis of SOCE in CD4^+^ T cells from *Orai1^fl/fl^Cd4^Cre^* mice transduced with Orai1α, Orai1β or EV. Cells were loaded with Indo-1 and stimulated with thapsigargin (TG) in Ca^2+^-free ringer solution followed by Ca^2+^ readdition (final concentration 1 mM Ca^2+^). SOCE was measured by flow cytometry. Shown are the slope, area under the curve (AUC) and peak of SOCE. Data are from 11 mice and four repeat experiments with one technical replicate per condition. (**B**) ImageStream analysis of endogenous NFAT1 localization using an anti-NFAT1 antibody. Representative images show brightfield (BF), CD4, Ametrine, NFAT1, and DAPI channels in Orai1-deficient CD4^+^ T cells transduced with Orai1α, Orai1β or EV and either left unstimulated or stimulated with 1 μM ionomycin to induce SOCE. (**C**) Quantification of NFAT1 nuclear translocation, expressed as similarity scores between NFAT1 and DAPI signals. (**D**) Cytokine production following PMA/ionomycin stimulation for 4 h. Representative flow cytometry plots of TNF, IFN-γ, and IL-2 in Orai1-deficient T cells transduced with either Orai1α, Orai1β, or EV gated on Ametrine^low^ cells. (**E**) Quantification of mean fluorescence intensities (MFI) of TNF, IFN-γ, and IL-2 in CD4^+^ T cells from D. Data are from 3 mice and two repeat experiments with one technical replicate per condition. Statistical analysis in panels (**A, E**) was conducted by ordinary one-way ANOVA multiple comparison. All results are expressed as means ± SEM. P values for the indicated comparisons are shown.

We next examined the ability of Orai1α and Orai1β expressed in *Orai1^CD4^*T cells to induce nuclear translocation of native NFAT1. Amt^low^ CD4^+^ T cells were enriched by flow cytometric cell sorting and stimulated with 1 μM ionomycin for 30 min or left untreated. Permeabilized cells were stained with anti-NFAT1 antibody and analyzed by ImageStream. Unstimulated T cells had predominantly cytosolic NFAT1, whereas stimulation induced robust nuclear translocation in *Orai1^CD4^* T cells transduced with Orai1 isoforms but not EV (**Fig. 5B-C**). Importantly, no differences regarding nuclear to cytoplasmic NFAT1 ratios were observed between Orai1α-and Orai1β-transduced CD4^+^ T cells. Because NFAT is essential for the transcriptional regulation of cytokine production in CD4^+^ T cells, we analyzed the levels of IL-2, TNF and IFN-γ in *Orai1^CD4^* T cells reconstituted with either Orai1α or Orai1β. CD4^+^ T cells were stimulated with phorbol 12-myristate 13-acetate (PMA) and ionomycin for 4 h followed by intracellular cytokine staining and analysis of Amt^low^ CD4^+^ T cells by flow cytometry. Orai1 reconstitution in *Orai1^CD4^*T cells significantly increased the expression levels of IL-2, TNF, and IFN-γ compared to mock-transduced cells and the frequencies of cytokine expressing cells (**Fig. 5D-E**). Again, no significant differences were observed between the ability of Orai1α or Orai1β to restore cytokine production in *Orai1^CD4^* T cells. Together these findings indicate that both Orai1 isoforms can effectively promote NFAT1 activation and cytokine gene expression in CD4^+^ T cells.

### Orai1α and Orai1β regulate similar transcriptional programs in primary CD4⁺ T cells

SOCE and NFAT regulate many other transcriptional programs in T cells besides cytokine expression including metabolism (14, 71, 72), CD4^+^ T cell differentiation (16, 72–76), anergy and exhaustion (77–79). To determine the broader transcriptional consequences of Orai1α and Orai1β signaling in CD4 T cells, we conducted RNA-sequencing of primary murine CD4⁺ T cells from *Orai1^CD4^* mice that were reconstituted with Orai1α, Orai1β, or EV. Cells were either left untreated or stimulated by anti-CD3/CD28 stimulation for 6h. We first confirmed isoform-specific expression of Orai1 by aligning sequencing reads to exons 1 and 2 of the *Orai1* gene (**Fig. 6A**). Principal component analysis (PCA) showed a clear separation between unstimulated and stimulated T cells (**Fig. 6B**). Within the stimulated samples, Orai1α-and Orai1β-expressing *Orai1^CD4^* CD4⁺ T cells clustered closely together and away from EV-transduced *Orai1^CD4^* T cells and WT controls. The analysis of differentially expressed genes (DEG) identified hundreds of genes that were either up-or downregulated in their expression in Orai1α- and Orai1β transduced *Orai1^CD4^* CD4⁺ T cells compared to EV controls (**Fig. 6C**). The total number of DEGs either up- or downregulated in Orai1α transduced cells was slightly larger than in cells expressing Orai1β (**Fig. 6C**). A correlation analysis of DEGs in T cells expressing either Orai1α or Orai1β, however, showed that log₂ fold-changes (FC) of gene expression were nearly identical between both isoforms (correlation coefficient r = 0.94); **Fig. 6D**), indicating that the differences in DEG counts in **Fig. 6C** are an artifact of statistical thresholding and not caused by distinct biological effects of both isoforms. Moreover, the analysis of the most upregulated genes in stimulated Orai1α transduced CD4⁺ T cells revealed that their expression levels were comparable to those in Orai1β transduced T cells (**Fig. 6E**). These genes include known NFAT-regulated cytokines such as IL-3, IL-10 and IFN-γ. Pathway interaction database (PID) analyses confirmed that both Orai1 isoforms regulated the same immune signaling pathways, including TCR, calcium and NFAT signaling, Polo-like kinase 1 (PLK1) signaling, and IL-2 signaling (**Fig. 6F**). Taken together, these results demonstrate that Orai1α and Orai1β drive nearly identical global transcriptional programs in T cells.

**Figure 6.**
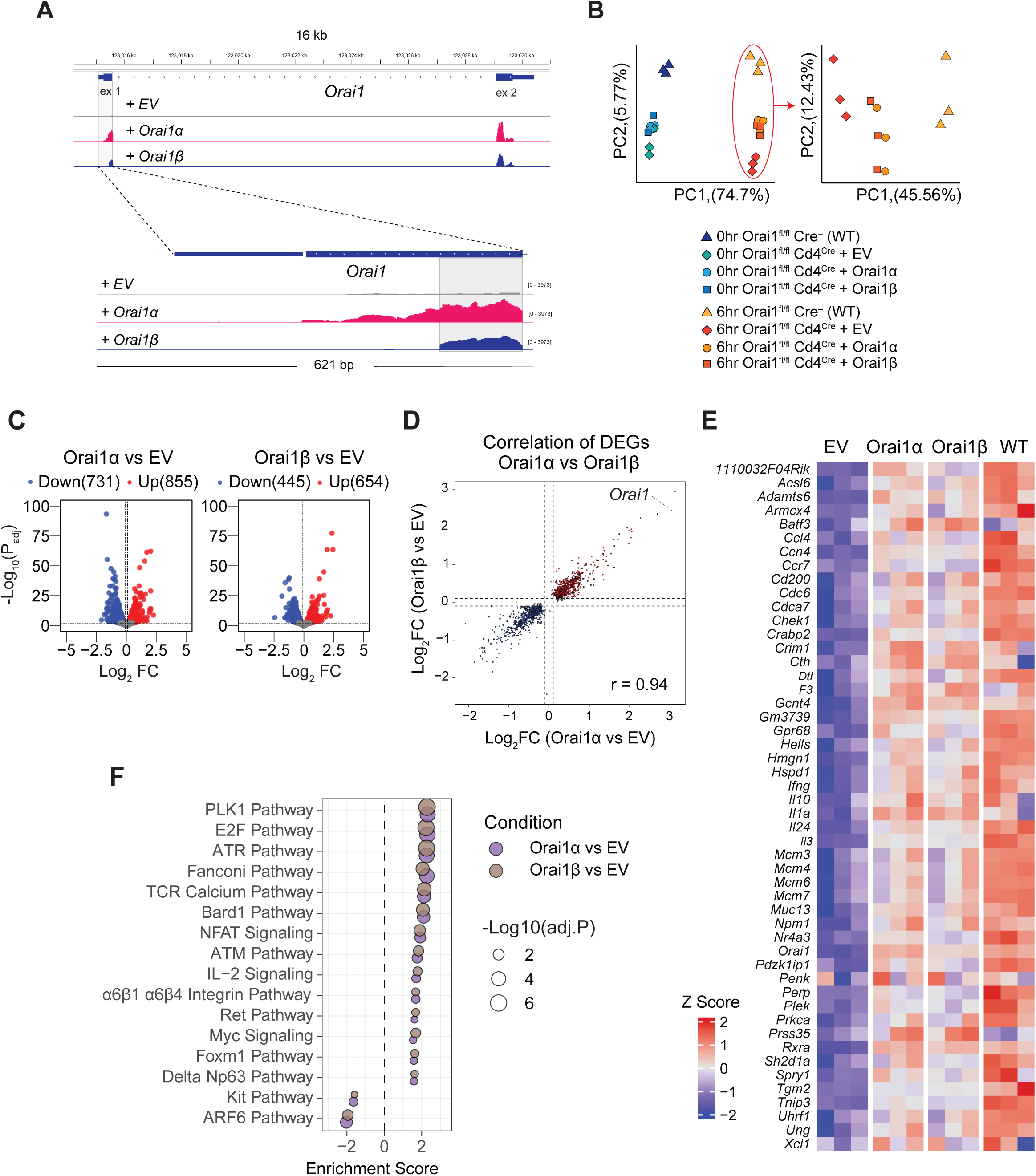
Orai1α and Orai1β orchestrate near-identical transcriptional programs in CD4⁺ T cells. Bulk RNA sequencing of CD4⁺ T cells from *Orai1^fl/fl^ Cd4^Cre^* mice were transduced with Orai1α, Orai1β, or empty vector (EV) and either left unstimulated (0h) or stimulated with anti-CD3/anti-CD28 for 6h. (**A**) BigWig tracks of ectopically expressed Orai1α and Orai1β mRNA mapped to the murine *Orai1* RefSeq (NM_175423). (**B**) Principal component analysis (PCA) of all RNA-seq samples (left) and of stimulated samples only (right). (**C–E**) Analysis of differentially expressed genes (DEGs). (**C**) Volcano plots comparing DEGs in CD4^+^ T cells expressing either Orai1α or Orai1β *vs* EV controls. (**D**) Dot plot of log₂ fold-changes (FC) in gene expression in Orai1α *vs* Orai1β-expressing CD4⁺ T cells. (**E**) Heatmap of the top 50 upregulated genes (ranked by significance) in stimulated Orai1α transduced CD4⁺ T cells compared to expression of the same genes in Orai1β and EV transduced or wildtype (WT) T cells. (**F**) Bubble plot of dysregulated pathways in Orai1α and Orai1β-expressing CD4⁺ T cells compared to EV controls using the Pathway Interaction Database (PID). Data in **A-F** are from three samples per condition. Statistical analyses in (**C**) were performed using the Wald test. DEGs in (**C, D**) were considered significant if the adjusted P value (P_adj_) was <0.01. The correlation coefficient (r) in (**D**) was calculated using Pearson’s method. Z scores in (**E**) were calculated using the average expression across all samples. The significance of pathways in (**F**) was calculated using a right-tailed Fisher’s exact test. Significance was adjusted using the Benjamini-Hochberg method.

### Homozygosity for a common frameshift mutation selectively abolishing Orai1α expression is not associated with the classical CRAC channelopathy phenotype

Homozygous null mutations in the *ORAI1* gene that either abolish protein expression or impair CRAC channel function in human patients cause immunodeficiency associated with recurrent severe infections early in life (41, 80). Patients also develop a characteristic set of non-immunological symptoms including amelogenesis imperfecta, anhidrosis and muscular hypotonia. Notably, all known *ORAI1* null mutations causing CRAC channelopathy are in residues shared by Orai1α and Orai1β and thus affect both isoforms (**Fig. 7A**). To evaluate the physiological role of Orai1α in humans, we used population genetic databases to identify and analyze genetic variants in the N terminus of Orai1 that exclusively affect Orai1α. We identified two frameshift mutations confined to this region (p.P43TfsX45 and p.P46SfsX42) that are predicted to selectively disrupt Orai1α expression without affecting Orai1β (**Fig. 7A**). Using immunoblotting of deglycosylated protein samples to distinguish between Orai1α and Orai1β, we confirmed that overexpression of the Orai1 p.P43TfsX45 variant in HEK293 Orai-TKO cells indeed resulted in the loss of Orai1α protein while preserving the expression of Orai1β, which is translated from M64 downstream of the mutation (**Fig. 7B**). Analysis of population genetic datasets including the Genome Aggregation Database (gnomAD), UK Biobank (UKBB), Qatar Biobank (QBB), and the All of Us (AoU) Research Program revealed that both N-terminal truncating variants are present at higher allele frequencies compared to verified disease-causing mutations affecting regions shared by both isoforms (**Fig. 7C**). Homozygous carriers of the Orai1α-specific truncating variants p.P43TfsX45 and p.P46SfsX42 were identified across multiple population cohorts, including 43 and 41 individuals in gnomAD, 2 and 0 individuals in QBB, and 8 and 8 individuals in AoU, respectively. By contrast, no homozygotes were identified in these databases for *ORAI1* loss-of-function mutations affecting regions shared by both Orai1 isoforms that are associated with CRAC channelopathy (41). These results suggest tolerance of mutations that selectively disrupt Orai1α while preserving Orai1β. To determine whether Orai1 p.P43TfsX45 and p.P46SfsX42 variants are associated with clinical phenotypes, we examined diagnostic codes associated with CRAC channelopathy due to *ORAI1* null mutations in UK Biobank (UKBB) (41). Individuals homozygous for the Orai1α-specific truncation variants did not exhibit an enrichment of disease codes for infectious diseases, ectodermal dysplasia or myopathy characteristic of CRAC channelopathy based on the analysis of UKBB and AoU databases (**Fig. 7D–E, Fig. S5A**). Only the prevalence of sweating disorders in homozygous carriers of *ORAI1* variants reached statistically significant levels in the AoU (but not UKBB) cohort. Further analysis of other diagnostic codes in individuals with Orai1α-specific truncation variants against the AoU cohort revealed a statistically significant increase in the prevalence of hyperhidrosis – albeit with incomplete penetrance – and a variety of other diagnostic codes (**Fig. S5B**). However, the significance of these findings is uncertain given the small number of affected individuals. Together, these findings indicate that selective disruption of Orai1α does not result in CRAC channelopathy disorder while the significance of its other disease associations remains unclear, at least based on available clinical genetic cohorts.

**Figure 7.**
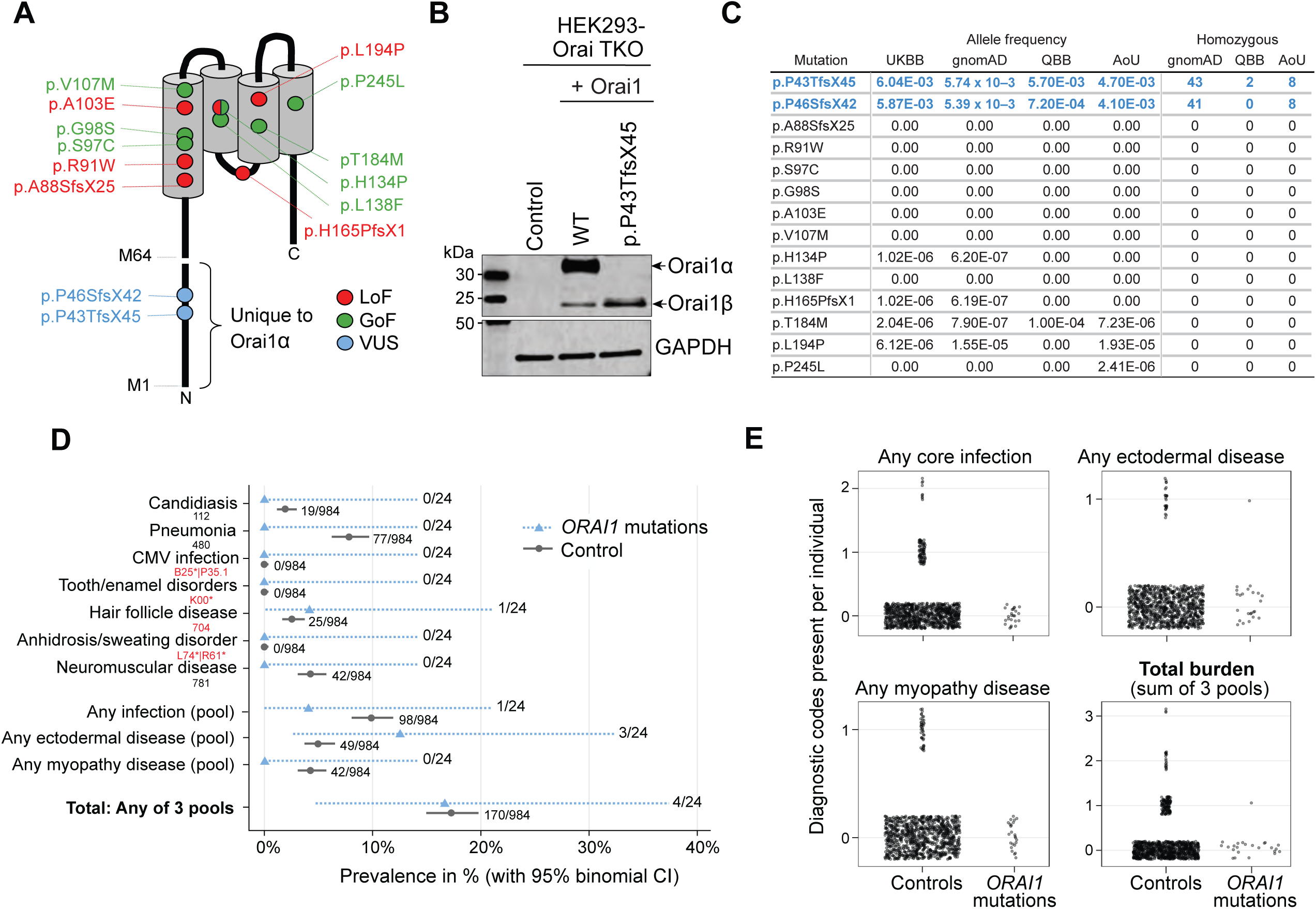
Human ORAI1 variants predicted to delete Orai1α protein expression lack clinical features associated with CRAC channelopathy syndrome. **(A)** Schematic representation of Orai1 channel topology and disease-associated loss-of-function (LoF, red) and gain-of-function (GoF, green) mutations reported in patients. Mutations are localized in regions shared by Orai1α and Orai1β. Also indicated are the locations of two frameshift mutations (blue) p.P43TfsX45 and p.P46SfsX42 in the Orai1α-specific N-terminal region. **(B)** Immunoblot analysis of WT Orai1 and the p.P43TfsX45 variant ectopically expressed in HEK293 Orai-TKO cells and detected with anti-Orai1 antibody. Indicated are the molecular weights of Orai1α and Orai1β isoforms. **(C)** Population genetic analysis of Orai1 variants in UK Biobank (UKBB), gnomAD, Qatar Biobank (QBB), and All of Us (AoU) databases. Allele frequencies (UKBB, gnomAD, QBB, AoU) and number of homozygous carriers (gnomAD, QBB, AoU) of each variant. **(D)** Prevalence of diagnostic codes in UKBB associated with symptoms of CRAC channelopathy syndrome (infection, ectodermal disease, myopathy) in 24 individuals homozygous s for the Orai1 p.P43TfsX45 and p.P46SfsX42 variants compared to 984 controls. **(E)** Distribution of disease burden (total and for specific diagnostic categories) in individuals homozygous for the Orai1 p.P43Tfs*45 and p.P46Sfs*42 variants compared to controls. Each dot represents one individual.

We identified individuals carrying the Orai1 p.P43TfsX45 variant in the Qatar Genome Project healthy population study (81). All individuals in this population study were recruited as healthy with no apparent clinical phenotype. We chose three individuals of the same gender but different ages who are homozygous, heterozygous and wildtype for the Orai1 p.P43TfsX45 variant, respectively. We used immunofluorescence to assess NFAT1 translocation both at baseline before stimulation and following store depletion with thapsigargin to activate SOCE. Stimulation induced significant nuclear translocation of NFAT1 that was more pronounced in both the heterozygous and homozygous donors compared to cells from the wild-type donor (**Fig. 8A-B**). This is consistent with increased NFAT1 translocation in HEK293 cells expressing Orai1β compared to Orai1α (**Fig. 2**). Interestingly we noted that even before stimulation the levels of NFAT1 in the nucleus were elevated in T cells from the homozygous and heterozygous individuals compared to the wild-type control (**Fig. 8B**). This suggests that even in the absence of maximal store depletion with thapsigargin, the loss of a single or both alleles of Orai1α induces higher baseline levels of NFAT1 translocation. Our studies in HEK293 cells argue that the extent of NFAT1 translocation depends on the magnitude of SOCE. We therefore measured SOCE in T cells from the three individuals who are homozygous, heterozygous and wildtype for the Orai1 p.P43TfsX45 variant. T cells from the individuals homozygous and heterozygous for Orai1 p.P43TfsX45 displayed elevated SOCE after thapsigargin stimulation compared to the wildtype control (**Fig. 8C-D**), consistent with results in HEK293 cells (**Fig. 3A**) and the enhanced fast CDI of Orai1α compared to Orai1β reported earlier (12). To test whether increased SOCE in the homozygous and heterozygous individuals is associated with altered STIM1 recruitment into puncta, a hallmark of CRAC channel activation (2), we immunostained endogenous STIM1 following store depletion and quantified the volume of STIM1 clusters after 3D reconstruction from a z stack of images at the single cell level. The results show no significant difference in the volume of STIM1 clusters in the three individuals arguing against differential STIM1 recruitment to Orai1α and Orai1β (**Fig. S6 A-B**). Together, these data demonstrate that an Orai1 mutation resulting in the expression of Orai1β alone does not impair, but rather enhances, SOCE and NFAT1 activation in primary human T cells.

**Figure 8.**
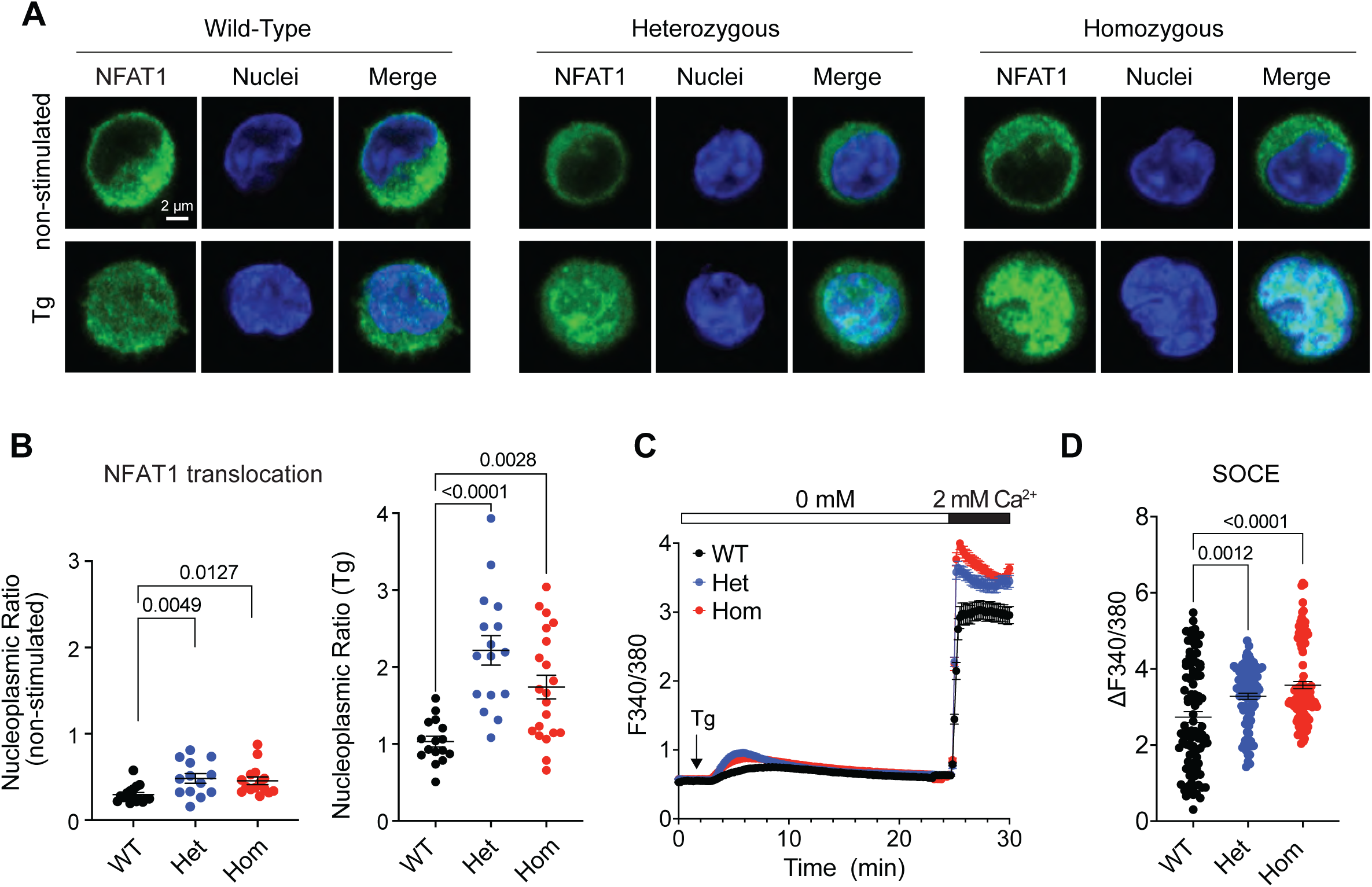
Functional studies of Orai1α deficient human T cells. **(A)** Confocal images illustrating the nuclear translocation of endogenous NFAT1 (green) induced by 1 μM Thapsigargin (Tg) in cells derived from individuals who are heterozygous (het) or homozygous (hom) for the Orai1α null variant (p.P43TfsX45) or wildtype (WT). **(B)** Quantification of endogenous NFAT1 translocation expressed as nuclear to cytoplasmic ratio before and after Tg stimulation. Each dot represents an individual cell. **(C)** SOCE recordings in T cells loaded with the Ca^2+^ dye Fura-2. Store depletion is induced in the absence of extracellular Ca^2+^ using Tg (1 μM), extracellular Ca^2+^ (2 mM) is added after 20 min to trigger Ca^2+^ influx through SOCE. **(D)** Quantification of peak SOCE amplitude after Ca^2+^ addition in T cells from the three genotypes. Data are expressed as mean ± SEM, statistical significance was assessed using one-way ANOVA with Dunnett’s post hoc test. For each individual, cells were from a single donor-derived culture. For Ca²⁺ imaging, recordings were obtained from 5 dishes for WT, 7 for the heterozygous individual, and 3 for the homozygous individual; data represent repeat recordings rather than independent biological replicates. Statistical comparisons were performed at the single-cell level.

## Discussion

Here we provide a comprehensive profiling of the coupling between native Orai channel homologues, SOCE and NFAT activation. While prior studies established that Orai1 is significantly more efficient than Orai3 at coupling to NFAT activation, they used overexpression models of STIM1, Orai1 and Orai3 and relied on heterologous expression of receptors for physiological stimulation (39, 40). Further, previous attempts to address the differences Orai1α and Orai1β coupling to SOCE and transcription in a native system were performed in Orai1 knockout HEK293 cells which nevertheless retained native expression of Orai2 and Orai3 isoforms (37). Here we used multiple CRISPR/Cas9-engineered HEK293 cell lines that express a single Orai homologue at native levels and are stimulated under conditions that mimic the range of physiological Ca²⁺ signaling events. This strategy allowed us to resolve quantitative differences in NFAT nuclear localization across Orai isoforms in response to relatively low and maximal levels of store depletion. Our data reveal that NFAT activation positively correlates with the amount of Ca²⁺ influx mediated by each Orai homologue and, as shown previously, has an inverse relationship with the extent of Ca²⁺-dependent fast inactivation reported for each homologue (12, 30, 37). We also compared the abilities of both Orai1 translational variants to mediate SOCE and activate NFAT. We found that in HEK293 cells Orai1β more potently induced NFAT1/4 nuclear translocation compared to Orai1α or the other Orai homologues, Orai2 and Orai3. Of note, the nuclear to cytoplasmic ratio in response to thapsigargin stimulation was greater for NFAT1 than NFAT4, consistent with the reported slower export kinetics of NFAT1(28).

We extended our analysis of Orai coupling to transcription in primary immune cells. We chose murine CD4⁺ T cells where Orai1 is indispensable for T cell activation, metabolic reprogramming, secretion of cytokines, and effector functions (14). In this physiological setting, we focused on the alternatively translated Orai1α and Orai1β isoforms and asked whether either could compensate for the loss of endogenous Orai1. Reconstitution with either isoform restored NFAT nuclear translocation, and cytokine production. Moreover, a global transcriptomic analysis of CD4⁺ T cells expressing either Orai1α or Orai1β revealed highly overlapping gene expression signatures that were enriched for canonical immune activation pathways, closely resembling wild-type CD4⁺ T cells. These results demonstrate that Orai1α and Orai1β are equally competent in driving the Ca²⁺-dependent transcriptional networks that underpin murine T cell effector function. This conclusion is further supported by human population genetic data and the analysis of human T cells from individuals carrying mutations that selectively abolish Orai1α protein expression. Individuals homozygous for variants that disrupt Orai1α without affecting Orai1β do not recapitulate the severe CRAC channelopathy phenotype associated with LOF mutations in regions shared by Orai1α and Orai1β (41). We cannot fully exclude that Orai1α and Orai1β may differentially regulate transcriptional programs in cell types other than T cells and that LOF mutations selectively impairing Orai1α expression cause disease phenotypes distinct from classical CRAC channelopathy syndrome.

While ectopically expressed Orai1α or Orai1β were equally able to promote SOCE, NFAT nuclear translocation and transcriptional responses in CD4*^+^*T cells from *Orai1^CD4^* mice, we observed a moderately greater ability of Orai1β compared to Orai1α to mediate SOCE and NFAT nuclear translocation in HEK293 cells and T cells from individuals homozygous or heterozygous for a Orai1α null variant. The reason for this discrepancy is not clear, but a possible explanation is that the ectopic expression levels of Orai1α and Orai1β in murine CD4^+^ T cells were sufficiently high to mask the subtle differences between both Orai1 isoforms as shown previously (12, 13). Increased SOCE and consequently NFAT nuclear translocation in HEK293 cells expressing low levels of Orai1β driven by a TK promoter construct and native Orai1β in human CD4^+^ T cells may be explained by previously described Orai1α-specific inhibitory mechanisms. Orai1β lacks two serine residues, S27 and S30, whose PKCβ mediated phosphorylation we had previously shown to inhibit SOCE and CRAC currents (36). SOCE is also inhibited by PKA-mediated phosphorylation of S34, which is lacking in Orai1β (37). In addition, we had demonstrated enhanced fast CDI of Orai1α compared to Orai1β (12). Because CDI determines how rapidly Orai-mediated currents decline, it can influence both the magnitude and kinetics of the resulting Ca²⁺ signal, and stronger inactivation of Orai1α has been associated with lower-frequency of agonist-evoked Ca²⁺ oscillations than Orai1β (37). We speculate that Orai1α- and Orai1β-specific CDI properties may shape the amplitude, duration and oscillatory frequency of SOCE. Whether CDI is altered in T cells of individuals with mutations in Orai1α but not murine T cells lacking Orai1α remains to be elucidated. Despite these differences, our data demonstrate that both Orai1α and Orai1β potently couple SOCE to NFAT activation and gene transcription. Moreover, our results identify the integrated strength of SOCE, which depends on either Orai1, Orai2 or Orai3 as the source of Ca^2+^ influx, as a major determinant of NFAT activation. A limitation of our study is that results are based on ectopic expression, albeit at near native levels, of individual Orai isoforms. It is likely that SOCE, NFAT nuclear translocation and gene expression are further shaped by different CRAC channel assemblies of heteromeric Orai subunits in different stoichiometries. These CRAC channel assemblies likely differ depending on the cell type considered and its activation state.

Our data using the Ca^2+^ chelators BAPTA and EGTA indicate that NFAT1 activation is mediated by localized Ca²⁺ signals near the CRAC channel pore, which is consistent with conclusions from previous studies of Orai1 channels (40) and more generally other Ca^2+^ channels such as voltage-gated Ca^2+^ channels (57, 66). How these localized Orai1-derived Ca^2+^ signals are converted into the activation of Ca^2+^-regulated transcription factors such as NFAT, however, is not well understood. A recent study showed that amino acids 39-59 of Orai1α form a domain that allows binding of AKAP79, the recruitment of calcineurin and thus, activation of NFAT near the channel pore (40). That is an attractive model to explain the coupling of localized Ca^2+^ signals to NFAT activation and gene transcription. While the study showed that Orai1β cannot effectively couple SOCE to NFAT activation (40), our findings, by contrast, demonstrate that Orai1β supports robust NFAT activation and transcription. The reasons for this discrepancy are currently not clear. Orai1α and Orai1β were reported to have different biophysical properties including distinct CDI properties as discussed above, pH dependence, protein stability, membrane organization, and structural or heteromeric assembly capabilities (4, 12, 36, 37, 82–84), which may allow each isoform to function more effectively under distinct signaling or environmental conditions. The conclusion that Orai1β is fully competent at coupling SOCE to NFAT activation and transcription is further supported by its evolutionary conservation from invertebrates such as *Drosophila Melanogaster* and *Caenorhabditis elegans* through vertebrates including zebrafish, frog, and chicken and all the way to mammals, including mouse, rat and human. By contrast, the Orai1α specific N-terminus is highly conserved only in mammals (**Fig. S7**) even though Ca^2+^-calcineurin-NFAT dependent transcription is an ancient signaling module that predates mammals (85–87). This phylogenetic evidence argues that Orai1β represents the ancestral isoform of Orai1 channels and is sufficient to effectively couple SOCE to transcription in most cells.

Future work will need to extend the physiological and pathophysiological roles of Orai1α and Orai1β in different cell types and organs in vivo. In immune cells, it will be important to define how Orai1α and Orai1β shape other immune functions such as T and B cell differentiation, effector functions, immunological memory and tolerance. Beyond immunity, clarifying whether Orai isoforms are differentially expressed in other cell types and tissues, or disease states will reveal whether changes in the relative proportions of Orai1α and Orai1β could bias transcriptional programs. While publicly available proteomics data indicate distinct expression patterns of Orai and STIM proteins across multiple human and mouse tissues and cell types, these datasets fail to distinguish between Orai1α and Orai1β as standard bottom-up proteomics predominantly detect peptides shared by both Orai1 proteoforms. The discovery of unique physiological or pathological functions of each Orai isoform could spur the development of pharmacological tools capable of selectively modulating each Orai isoform for therapeutic interventions in disease.

## Methods

### Cell culture and transfection

HEK293 cells obtained from ATCC (Catalog # CRL-1573) were grown in Dulbecco’s modified Eagle’s medium (DMEM) containing high glucose (4.5 g/L), supplemented with 1% Antibiotic– Antimycotic solution and 10% heat-inactivated fetal bovine serum. The cells were incubated in a humidified CO2 incubator at 37 °C under standard cell culture conditions (5% CO_2_, 95% air).

Transfections were conducted using lipofectamine 2000 with a 1:3 Lipofectamine: cDNA ratio. 75-100K cells were seeded onto 25 mm glass coverslips and incubated for 24 h prior to experiments. During transfection, the Antibiotic–Antimycotic solution was removed from the media. For experiments performed in Orai double knockout HEK293 cells (DKO), 1.0 μg of NFAT-GFP plasmids were used for transfection. For experiments performed in Orai triple knockout HEK293 cells (TKO), 1.0 μg of NFAT-GFP plasmid together with either 0.3–0.5 μg of TK-Orai1α, TK-Orai1β, TK-Orai2 or TK-Orai3 plasmids were used. Transfections were confirmed by recording the corresponding fluorescence, calcium imaging and immunoblotting or qPCRs.

### Genetic knockout of *ORAI* genes in HEK293 cells using CRISPR/Cas9

The creation of Orai-KO HEK293 cell lines was performed in two steps. The first step involved using a single *ORAI*-specific gRNA to generate insertions/deletions in the *ORAI* gene through transfection of corresponding *ORAI1* or *ORAI3* targeting vector followed by puromycin selection, colony screening by Guide-it Mutation Detection Kit, and Sanger sequencing for genetic knockout confirmation. This step was used to generate single knockouts (SKO), Orai1-SKO and Orai3-SKO clones. The second step was to achieve double knockout (DKO) and subsequently triple knockout (TKO) in the SKO clones. This involved subcloning of two *ORAI2* or *ORAI3* specific gRNAs flanking the entire gene into fluorescent vectors (pSpCas9(BB)-2A-GFP and pU6-(BbsI)-CBh-Cas9-T2AmCherry: Addgene plasmid #48138, and plasmid #64324) and sorting single cells with high GFP and mCherry expression into 96-well plates. Positive clones were identified through screening with specific PCR primers distinguishing the WT and Knockouts and confirmed by Sanger sequencing. All gRNA sequences used can be found in our previously published study(32).

### Generation of Orai1-knockout MCF-7 cells (CRISPR/Cas9)

MCF-7 cells (ATCC, HTB-22) were engineered to delete ORAI1 using CRISPR/Cas9 gene editing (29). A gRNA targeting human *ORAI1* (GTTGCTCACCGCCTCGATGT) was cloned into lentiCRISPR v2 (Addgene) using the following oligonucleotides: PX11 (5′- CACCGGTTGCTCACCGCCTCGATGT-3′) and PX12 (5′-AAACACATCGAGGCGGTGAGCAACC-3′). Cells were transfected with the gRNA–Cas9 construct, puromycin-selected for 5 days, and plated at single-cell density in 96-well plates. Expanded clones were genotyped by PCR across the cut site (primers PX39 and PX11) and re-screened with flanking primers (PX39 and PX3). Amplicons were cloned and sequenced to confirm indels in all *ORAI1* alleles leading to frameshift and premature stop codons. Validated Orai1-KO clones were further confirmed by loss of Orai1-dependent SOCE and used for Ca²⁺ imaging and NFAT1 activation assays.

### Constructs and transfections

To produce a vector that exclusively expresses the full-length, long Orai1α isoform, a wild-type ORAI1-CFP cDNA clone (Addgene plasmid #19757)was utilized. This involved replacing the weak native ORAI1 Kozak sequence located upstream of the first ATG start site (TGCTCCATG) with a stronger Kozak sequence (GCCACCATG). This resulted in the translational machinery being forced to start almost exclusively at the first ATG. Conversely, to create a vector that exclusively expresses the short Orai1β isoform, the first ATG start site of ORAI1-CFP cDNA was mutated to GCG (alanine) or ATC (Isoleucine), which left only the start site at methionine-64. Further details of how Orai1α and Orai1β vectors were constructed have been described previously (37). Constructs were validated by sequencing, and expression in cells was validated by fluorescence imaging and Western blotting (**Fig. S1A-E**) and functionally by their ability to rescue SOCE in HEK293 TKO cells. The florescence tags in both constructs were swapped to either YFP, m-Cherry, or Ametrine instead of CFP based on the experimental needs. TK-driven Orai1/2/3 constructs, and gRNA were generated using primers compatible with the in-fusion cloning system. The resulting plasmids were confirmed by sequencing and by PCR using selective primers to ensure proper cDNA insertion or mutation. NFAT1/4-GFP plasmids are from Addgene (Plasmid #11107 and **#**21664).

### Assessment of expression of YFP/CFP-tagged Orai isoforms in Orai-TKO HEK293 cells

The expression of Orai1-YFP in HEK293 cells was confirmed by capturing cell fluorescence using a Leica DMI8 confocal microscope with a Plan-Apochromat 40×/1.4 oilDIC M27 objective at 1× zoom. The 488-nm laser line was used to excite YFP, and fluorescence emission was detected between 491 nm − 695 nm with a 9-nm bandwidth, using digital photon counting mode and a pinhole of 180 μm with 8-line averaging. Expression was additionally quantified in 6-well plates using a BioTek Synergy Neo2 plate reader in fluorescence area-scan mode: YFP-tagged constructs were read at 513/20–488/20 nm excitation and 527/20 nm emission, and CFP-tagged constructs were read at 433/20 nm excitation and 475/20 nm emission, normalized to Fura-2 fluorescence (380/20 nm excitation, 510/20 nm emission).

### Western blot analysis

The cells were rinsed with ice-cold phosphate-buffered saline (PBS) and then lysed in RIPA buffer supplemented with protease and phosphatase inhibitors at 4 °C for 30 min. The lysates were centrifuged at 14,000 RPM for 15 min at 4 °C, and the clarified supernatant was collected for protein concentration measurement using the Pierce BCA assay. To perform deglycosylation either

Protein Deglycosylation Mix II O-Glycosidase kit (NEB #P0733) or N-Glycosidase F (Millipore Sigma) were used. For the Protein Deglycosylation Mix II the manufacturer instructions were followed, and for N-Glycosidase F (Millipore Sigma) 5 μl of N-Glycosidase were added to 50 μg of protein lysate and incubated in a 37°C water bath overnight. Afterward, 30-60 μg of protein lysate was loaded into a 4-12% Bis-Tris gel. The deglycosylated protein extracts were loaded on NuPAGE 4–12% Bis-Tris precast protein gels and subjected to electrophoresis. The proteins were then transferred to polyvinylidene difluoride (PVDF) membranes overnight at 4 °C. Subsequently, the membranes were blocked with LI-COR Tris-buffered saline (TBS) buffer and incubated overnight at 4 °C with anti-Orai1 polyclonal antibody (Cat # O8264, Sigma-Aldrich). The probed membranes were then washed with Tris-buffered saline / Tween-20 (TBST) and incubated with secondary antibodies for 1 h at room temperature with shaking. Finally, after washing with TBST the blots were imaged using the LI-COR Odyssey imaging system and quantified by Image Studio software (LI-COR).

### Ca^2+^measurements of HEK293 cells

Fluorescence measurements were performed in HEK293 cells loaded with the ratiometric calcium-sensitive dye, Fura-2 AM. Briefly, Type I collagen from calf skin (Sigma-Aldrich, Cat. No. C8919) was used to coat 25 mm glass coverslips at a concentration of 30% in a 30% ethanol solution for 30 minutes prior to cell seeding. Day 0: 48 hours before imaging, the cells were seeded on the coated coverslips at a density of 1×10^5^ cells per coverslip and allowed to settle for 24 h in the incubator. Day 1: transfections were performed as mentioned above. Day 2: coverslips were mounted into an Attofluor cell chamber (Thermo Fisher). The cells were then incubated with 2 μM Fura-2 AM (Thermo Fisher) in HEPES-buffered salt solution (HBSS) containing 140 mM NaCl, 4.7 mM KCl, 1.13 mM MgCl_2_, 2.0 mM CaCl_2_, 10 mM glucose, and 10 mM HEPES (pH 7.4) for 30 minutes at room temperature, washed four times with HBSS, and imaged using a Leica DMi8 fluorescence microscope with a 20X Fluor objective. Fura-2 was excited at 340 and 380 nm using a fast shutter wheel (Sutter Instruments), and emissions were captured at a wavelength of 510 nm with a Hamamatsu Flash 4 camera. The ratio of fluorescence emission upon alternate excitation at 340 nm and 380 nm (F340/F380) was analyzed using regions of interest drawn around the perimeters of 10-20 cells per coverslip with the Leica Application Suite X (LAS X) software. The mean ± SEM traces resulting from this analysis were represented.

### NFAT-GFP nuclear translocation assays in HEK293 cells

After HEK293 cell transfections with NFAT1-GFP or NFAT4-GFP as described above, GFP fluorescence in the cytoplasm and nucleus was collected on a Leica DMi8 microscope with a 20 × objective and a Hamamatsu Flash 4 camera, with Leica Application Suite X software used for image acquisition and analysis. GFP was excited with a 488 nm fast wheel filter, and its emission was selectively captured through a 510 nm GFP filter cube. Images were acquired every 15 s. For each cell, nuclear and cytoplasmic regions of interest (ROIs) were manually defined based on cell morphology and GFP fluorescence distribution, without nuclear counterstaining. NFAT nuclear translocation was quantified as ratio of nuclear to cytoplasmic GFP fluorescence intensity measured at 510 nm emission(F510) for 1 min prior to stimulation (baseline) and in response to 10 μM CCh or 2 μM thapsigargin (Tg) as previously described (31, 32, 37). Background fluorescence was subtracted from each ROI, and the fluorescence of each cell was normalized to its baseline fluorescence value to minimize cell-to-cell variability in expression levels.

### Ca^2+^ chelation experiments

To distinguish between effects of local and global Ca²⁺ signaling, HEK293 Orai-TKO cells transfected with either Orai1α or Orai1β were pre-incubated with either 20 μM EGTA-AM (Thermo Fisher Scientific, Cat. E1219) or 20 μM BAPTA-AM (Thermo Fisher Scientific, Cat. E1205) for 25 min at 37 °C prior to imaging. Following chelator loading, cells were washed and loaded with 2 μM Fura-2-AM as described above for SOCE measurements. For NFAT1 nuclear translocation experiments, cells expressing NFAT1-GFP were preloaded with EGTA-AM or BAPTA-before 2 μM thapsigargin stimulation. SOCE was quantified with Fura-2 340/380 ratio measurements, while NFAT activation was measured as the nuclear/cytoplasmic fluorescence ratio as described above.

### Mice

*Orai1^fl/fl^Cd4^Cre^* mice were generated by crossing *Orai1^fl/fl^* mice (64) to *Cd4^Cre^*mice (RRID:IMSR_JAX:017336). Sex-matched male and female mice were used between 10 and 12 weeks of age. Mice were maintained under specific pathogen–free conditions. All experiments were conducted in accordance with protocols approved by the Institutional Animal Care and Use Committee at New York University Grossman School of Medicine.

### Murine T cell culture

CD4^+^ T cells from *Orai1^fl/fl^Cd4^Cre^* mice were isolated from spleens using EasySep™ Mouse CD4^+^ T Cell Isolation Kit (Stemcell Technologies, Cat# 19852A) following the vendor’s instructions. CD4^+^ T cells were cultured in complete RPMI-1640 medium (Corning, 10-040-CV) containing 10% FBS, 2 mM L-glutamine, 50 U/ml penicillin-streptomycin, and 5.5 μM β-mercaptoethanol. For retroviral transduction, CD4^+^ T cells were seeded at a density of 1×10^6^/mL onto 12-well plates coated with 20 μg/mL rabbit-anti-hamster IgG (Thermo Scientific, Cat. A18891). Cells were then stimulated with anti-CD3ε (1 μg/mL, BioXcell, Cat. BE0001-1, Clone 145-2C11) and anti-CD28 (1 μg/mL, BioXcell, Cat. BE0015-1, Clone 37.51) antibodies in the presence of 5 μg/mL anti-IFN-*γ* (BioXcell, Cat. BE0055, Clone. XMG1.2) and anti-IL-4 (BioXcell, Cat. BE0045, Clone. 11B11) antibodies. Twenty-four hours post activation, T cells were transduced with retroviral supernatant and 8 μg/ml Polybrene (Sigma-Aldrich, Cat. 107689) by spin-transduction (2.500 rpm, 90 min, 32°C). Retroviral supernatant from T cells was removed 30 min after spin infection and replaced with fresh complete RPMI-1640 media. T cells were detached 24 h later and split into 6-well plate with 5 mL of complete RPMI-1640 media containing 20 U/mL recombinant human (rh) interleukin (IL)-2 (PeproTech, Cat. 200-02) and IL-12 (20 ng/mL, PeproTech, Cat. 210-12) to expand cells for 2 more days before analysis. Retroviral supernatant was produced in the Platinum-E retroviral packaging cell line (88). Platinum-E cells were transfected by GeneJet (Fisher, Cat. FERK0481) with retroviral expression plasmids pMSCV-Orai1α-IRES-Ametrine, pMSCV-Orai1β-IRES-Ametrine or pMSCV-IRES-Ametrine (empty vector, EV) with the ecotropic packaging vector pCL-Eco. Retroviral supernatants were collected 36 and 60h after transfection.

### Murine CD4^+^ T differentiation assays for Orai1α/β expression

Naive CD4^+^CD25^−^CD44^low^ T cells were purified from spleens and lymph nodes (cervical, inguinal, brachial, and axillary) of wild type C57BL/6 mice using a CD4^+^ T cell isolation kit (Miltenyi Biotech) according to the manufacturer’s protocol. Purified cells were activated at a density of 1 × 10^5^ cells per well in flat-bottom plates with either plate-coated anti-CD3 (4 μg/ml) and soluble anti-CD28 (2 μg/ml) for Th17 cells or anti-CD3 (2 μg/ml) and anti-CD28 (2 μg/ml) for Th1, Th2, or Treg cells. CD4^+^ T cell differentiation conditions were as follows: nonpathogenic (np) Th17 cells: 2.5 ng/ml rhTGF-β1 (BioLegend, Cat. 781802) plus 40 ng/ml rmIL-6 (BioLegend, Cat. 575702); pathogenic (p) Th17 cells: 40 ng/ml rmIL-6, 40 ng/ml rmIL-23 (BioLegend, Cat. 589002) plus 40 ng/ml rmIL-1β (BioLegend, Cat. 575102); Th1 cells: 20 ng/ml rmIL-12 plus 20 ng/ml rmIL-2; Th2: 20 ng/ml rmIL4, 20 ng/ml rmIL-2 plus 10 μg/ml anti-IFNγ; and iTreg polarization: 3 ng/ml rhTGF-β1 plus 20 ng/ml rmIL-2.

### Intracellular Ca^2+^ measurements of murine T cells

The flow cytometric analysis of intracellular Ca^2+^ levels in mouse CD4^+^ T cells was described previously (89). Briefly, retrovirally transduced CD4^+^ T cells from *Orai1^fl/fl^Cd4^Cre^* mice were first stained with either PE-Cy7 conjugated anti-CD4 (Biolegend, Cat. 100528) or APC-Cy7 conjugated anti-CD4 antibodies (Biolegend, Cat. 100414). Cells were then washed, combined, and loaded with 2 μM Indo-1 in fresh complete RPMI-1640 medium for 15 min. After Indo-1 loading, cells were washed and resuspended in 510 μL of Ca²⁺-free Ringer buffer. Intracellular Ca²⁺ levels were measured using an LSR Fortessa flow cytometer. Baseline fluorescence was recorded for 100 s followed by addition of 1 μM Thapsigargin (Sigma-Aldrich, Cat. No. 586005) to induce ER Ca²⁺ store depletion. At 400 s, 240 μL of Ca²⁺-containing Ringer buffer was added to the cell suspension (final Ca^2+^ concentration 1 mM) to induce SOCE. Data acquisition continued for a total of 600 s. Fluorescence signals were collected in BUV396 and BUV496 channels. Intracellular Ca²⁺ levels were quantified as the fluorescence emission ratio of BUV396 to BUV496 channels. The slope, area under curve (AUC) and peak of Ca^2+^ influx were calculated using Prism Software (GraphPad).

### ImageStream analysis of NFAT translocation in T cells

Nuclear translocation of NFAT1 in primary murine CD4⁺ T cells was assessed by imaging flow cytometry. CD4⁺ T cells from *Orai1^fl/fl^Cd4^Cre^* were retrovirally transduced with empty vector (EV), Orai1α or Orai1β, and enriched for Ametrine^low^ cells by fluorescence-activated cell sorting (FACS) as described above. 3 × 10^6^ cells per condition were either left unstimulated or stimulated with 1 µM ionomycin for 30 minutes at room temperature. Cells were fixed, permeabilized and incubated with an anti-NFAT1 rabbit monoclonal antibody conjugated with Alexa Fluor 488 (clone D43B1; Cell Signaling Technology, Cat. #14324) at a 1:100 dilution in 100 µL of Flow Staining Buffer for 30 minutes at room temperature in the dark. Cells were washed twice with 1× Flow Cytometry Perm Buffer and once with Flow Staining Buffer, then resuspended in 40 µL of 1× PBS. DAPI (10 µL of 1.25 µg/mL in 1× PBS) was added immediately before acquisition to a final concentration of 0.25 µg/mL to stain nuclei.

Images were acquired on an Amnis ImageStream X Mk II imaging cytometer. Brightfield images were obtained with channels 1 and 9 (430–480 nm); DAPI nuclear signal was collected in channel 7 (excitation 405 nm, emission 430–505 nm); and NFAT1-AF488 signal was collected in channel 2 (excitation 488 nm, emission 528–565 nm). Excitation laser power was adjusted prior to acquisition to prevent pixel saturation. Data were analyzed using IDEAS software version 6.2 (Amnis Corporation) using the built-in transcription factor translocation guided analysis workflow. Gating was applied to select single, in-focus cells that were double-positive for DAPI and NFAT1 staining. Nuclear translocation was quantified using the similarity score, a log-transformed Pearson correlation coefficient between the NFAT1 signal intensity and the DAPI-defined nuclear mask. Raw similarity scores were exported to CSV files for statistical analysis.

### Flow cytometry and cytokine production of murine T cells

Intracellular cytokine staining of murine CD4^+^ T cells was performed as previously described (74). Briefly, CD4^+^ T cells were stimulated with 1 μM ionomycin (SigmaAldrich, Cat. 407952) and 20 nM phorbol 12-myristate 13-acetate (PMA; Calbiochem, Cat. 524400) for 4 hours in the presence of 5 μM Brefeldin A (eBioscience, 00-4506-51). Cell surface marker staining was conducted first, cells were then fixed with an intracellular fixation kit (Invitrogen, 88-8824-00) for 20 min, permeabilized with Permeabilization Buffer (eBioscience, 00-8333-56) and incubated with anti-cytokine antibody cocktails (listed in **Supplementary Table S1**) for 30 min. Fluorescence signals were acquired using an LSR Fortessa flow cytometer and data were analyzed using FlowJo software (FlowJo v.10.8.1).

### Bulk RNA-sequencing

CD4^+^ T cells from *Orai1^fl/fl^Cd4^Cre^* mice that had been retrovirally transduced with empty vector (EV), Orai1α or Orai1β were stimulated with anti-CD3ε (1ug/mL) anti-CD28 (1 μg/mL) on 20 μg/mL rabbit-anti-hamster IgG (Thermo Scientific, Cat. A18891) coated 12-well plates for 6 hours and enriched for Ametrine^low^ (lower 25%ile) cells by flow cytometric cell sorting using a FACSAria II instrument (BD Biosciences). RNA was purified using the RNeasy Micro RNA Isolation Kit (Qiagen, Cat no. 74104) and RNA-Seq libraries were prepared using the Clonetech SMART-Seq HT Kit (Takara, Cat no. 634792). The amplified libraries were purified using AMPure beads (Beckman Coulter, Cat no. A63882), quantified using a Qubit 2.0 fluorometer (Life Technologies), and visualized on an Agilent Tapestation 2200. The libraries were pooled equimolarly, loaded onto on a NovaSeq6000 system (Illumina) using a SP 100 Cycle Flow Cell v1.5 to generate 100 base pair paired-end reads. FASTQ files were generated using the bcl2fastq2 conversion software for each sample (v2.20). Raw FASTQ files were assembled using Seq-N-Slide pipeline (DOI: 10.5281/zenodo.5550459). For gene expression analysis, reads were trimmed using trimmomatic (v0.36) and then aligned to the mouse reference genome (GRCm38.85 / mm10) using STAR aligner (v.2.5.0c). Feature count matrices were generated with the featureCounts function of the subread package (v2.0.3). Differential expression was performed in an R statistical environment with the DESeq2 (v1.50.2) package. Gene set enrichment analysis was performed using clusterprofileR(v4.18.4) using gene sets acquired from the molecular signature database.

### Population genetic analysis of ORAI1 variants

Allele frequencies and homozygous carrier counts for ORAI1 variants were obtained from the UK Biobank (UKBB), Genome Aggregation Database (gnomAD), Qatar Biobank (QBB), and the All of Us (AoU) Research Program (90). Variants were classified according to whether they selectively affected the Orai1α-specific N-terminus or regions shared by both Orai1α and Orai1β. Allele frequencies and numbers of homozygous carriers are summarized in Figure 7C. Approximately 40 controls were matched to each individual with homozygous ORAI1 variants. Because UKBBparticipants are socially and genetically heterogeneous, the control-matching algorithm optimized matches using characteristics including age, sex, healthcare utilization metrics, genetic ancestry, and genome-wide genetic principal components (PCs). This approach aimed to ensure that observed differences between cases and controls were attributable, as much as possible, to the variant of interest. A similar number of matched controls (∼40 controls per individual with homozygous ORAI1 variants) were used for the QBB analysis.

### QBB human study population

QBB is a prospective, population-based cohort study established in 2012 in Qatar (91). Participants are adults who are either Qatari nationals or have resided in Qatar for at least 15 years. All participants provided written informed consent before inclusion. The study was approved by the Hamad Medical Corporation Ethics Committee and the QBB institutional review board.

### Whole-genome sequencing

Genomic DNA was extracted from peripheral blood using the automated QIASymphony SP instrument according to the manufacturer’s instructions (Qiagen, Germany). DNA concentrations were quantified using the Quant-iT dsDNA Assay (Invitrogen, USA) on a FlexStation 3 (Molecular Devices, USA). Whole-genome sequencing was carried out using 2 × 150 bp paired-end reads on a HiSeq X Ten sequencer (Illumina, USA) at the Sidra Clinical Genomics Laboratory Sequencing Facility, achieving a mean coverage depth of 30×. Read-level quality control was assessed with FastQC (v0.11.2). Reads were aligned to the GRCh37 (hs37d5) reference genome using bwa.kit (v0.7.12). Post-alignment quality control of mapped reads was performed with Picard (v1.117, CollectWgsMetrics).

Variant calling followed GATK 3.4 best practices. Indel realignment and base quality score recalibration were applied to the initial BAM files, after which HaplotypeCaller was run on each sample to generate per-sample genomic variant call files (gVCFs). Joint variant calling was then performed across all samples: GenomicsDB was used to combine samples by genomic region, and GenotypeGVCFs was subsequently applied to each region. Variant quality score recalibration (VQSR) was performed separately for SNVs and indels. In total, 134 million variants passed the VQSR filter across the 14,669 QBB individuals (91). No individuals were excluded for high missingness (>0.1), as assessed using Plink (v2.0) (91). Approximately five million variants were removed due to a low call rate (<0.05).

### Isolation of human T cells

Peripheral blood mononuclear cells (PBMCs) were provided by Qatar Biobank (QBB) and experimentation was performed following approved QBB IRB protocol number E-2023-QF-QBB-RES-ACC-00153-0245. To activate and expand human T cells, thawed PBMCs were resuspended in complete Advanced RPMI 1640 medium supplemented with GlutaMAX (Gibco) and 10% fetal bovine serum (FBS). Cells were stimulated with 1 μg/ml purified mouse anti-human CD3 (clone HIT3a; BD Pharmingen) and anti-human CD28 (clone CD28.2; BD Pharmingen) antibodies and incubated at 37 °C in a humidified incubator with 5% CO₂ for 48 hours. Following activation, cells were maintained in culture in the presence of 30 U/ml recombinant human IL-2 (hIL-2; PeproTech). Fresh IL-2-containing medium was added every two days starting on day 3 post-activation, and T cells were expanded for up to 21 days.

### Intracellular Ca^2+^ imaging of human T cells

Ca^2+^ imaging was performed using a PTI Easy Ratio Pro system (software version 1.6.1.0.101; Horiba Scientific) composed of a DeltaRAMX monochromator and a CoolSnapHQ^2^ camera attached to an Olympus IX71 inverted microscope fitted with a 20x/0.75 lens. The standard saline contained (in mM) 145 NaCl, 5 KCl, 2 CaCl_2_, 1 MgCl_2_, 10 Glucose, 10 HEPES, pH 7.2, for Ca^2+^-free experiments, the Ca^2+^ was exchanged equimolarly with Mg^2+^. For Ca^2+^ imaging the cells were first plated for 30 minutes on CellTak (Corning) coated glass coverslips and were then loaded for 30 min with 2 µM Fura2-AM in a Ca^2+^-containing media at room temperature and washed three times in Ca^2+^-free media prior to the experiments. Cells were excited at 340 nm and 380 nm for 100 msec at a 0.1 Hz frame rate and the ratio of the fluorescence intensity at 340/380 measured. Tg (1 µM) was applied for 20 minutes (enough for the Ca^2+^ baseline to recover) in a Ca^2+^-free media. The extracellular Ca^2+^ concentration was then raised to 2 mM through direct injection of Ca^2+^ into the dish. The Ca^2+^ traces were analyzed using Clampfit (11.4.3, Molecular Devices).

### Immunocytochemistry of human T cells

T cells were plated as indicated for Ca^2+^ imaging and treated with thapsigargin (Tg, 1 µM) for 10 min before fixation for 10 min with 4% paraformaldehyde. Cells were then permeabilized with 0.1 % Triton X100 for 10 min and saturated for 1h using a mix of 10% goat serum and 1% bovine serum albumin. Incubation in the primary antibody for NFAT1 (5861S, Cell Signaling) was performed overnight at 4°C, secondary antibody was anti-rabbit coupled Ax488 (Molecular Probes) and used at a 1:2000 dilution at room temperature for 2 hours, staining of the nuclei was performed using Hoechst (8 µM). Confocal images were acquired using a Zeiss LSM880 confocal microscope (63x/1.4, 1 Airy Unit) and the following parameters (λ_ex_=405, λ_em_=410-495 for Hoechst, λ_ex_=488, λ_em_=495-630 for Ax488, 1AU). Z-stacks were recorded at 1 µm intervals. Analysis was performed using FIJI (92). Fixed T cells (see main manuscript methods) were incubated with a STIM1 primary antibody overnight at 4°C (4916S, Cell signaling,) and for 2 hours at room temperature with an Ax555 coupled secondary antibody (Molecular Probes) and counterstained with Hoechst to stain the nuclei. High resolution images were acquired using the Airyscan detector of a Zeiss LSM880 confocal microscope using the super-resolution mode (SR) and default image processing parameters. The 405 and 561 laser lines were used to acquire alternating z-stacks typically 0.18 µm intervals. Clusters identification was performed using the 3D Spot Segmentation plugin of FIJI using the watersheding option and default parameters. Spot seeding threshold was defined as the intensity in the middle of the weakest well defined cluster. Cluster volume measurement was performed using the built-in 3D object counter of FIJI. Outliers were removed from the data using Graphpad Prism “identify outliers” function. Because of the limited availability of these clinical specimens, all experiments for each individual were performed from a single donor-derived culture, and independent biological replication was not available. For NFAT1 translocation, a single set of cells was analyzed for each condition. For Ca²⁺ imaging, cells were plated directly from the suspension culture, and recordings were repeated across dishes to obtain comparable total cell numbers across conditions. In total, 5 dishes were recorded for WT, 7 for the heterozygous individual, and 3 for the homozygous individual. Statistical comparisons were performed at the single-cell level.

### Statistics

GraphPad Prism 8 software was used to conduct statistical analyses. For imaging experiments in which multiple cells were recorded within an independent experiment, cell-level measurements were averaged within each experiment, and the resulting experiment-level means were used as the unit of statistical analysis. Individual-cell measurements are shown for visualization where indicated. All bioinformatics analyses were performed in an R statistical environment (v4.2). The Student’s *t*-test or Welch’s *t*-test, as indicated, was used to compare two groups, while one-way analysis of variance (ANOVA) with Dunnett’s multiple-comparisons test was used to compare more than two groups. Results are expressed as mean ± SEM. Exact *P* values are shown where indicated. For the human T-cell experiments in Figure 8, independent biological replication was not available because cells from each individual were derived from a single donor-derived culture. Statistical comparisons for these experiments were therefore performed at the single-cell level, as described in the corresponding Methods and figure legend.

## Data availability

The RNA-seq data generated by this study are publicly available through the GeneExpression Omnibus under accession number GSE334106.

## Acknowledgments

This research has been conducted using the UK Biobank Resource under Application Number 105564.

## Author Contributions

A.E.A., K.M, S.F., and M.T. designed research; A.E.A., Y.W., J.C.B., I.P., B.S., M.M., M.J., A.T.E., S.C.-R., R.C., A.B., F.Y., D.S.P., S.Y., and P.X. performed research; A.C.S., F.F.S., N.H., W.F.H., D.B.B., K.M., S.F., and M.T. contributed new reagents, analytic tools, or clinical/genetic resources; A.E.A., Y.W., J.C.B., I.P., B.S., M.M., M.J., R.C., A.B., F.Y., S.Y., D.B.B., K.M., S.F., and M.T. analyzed data; and A.E.A., K.M., S.F., and M.T. wrote the paper with input from all authors.

## Supplementary Figure legends

**Figure S1.**
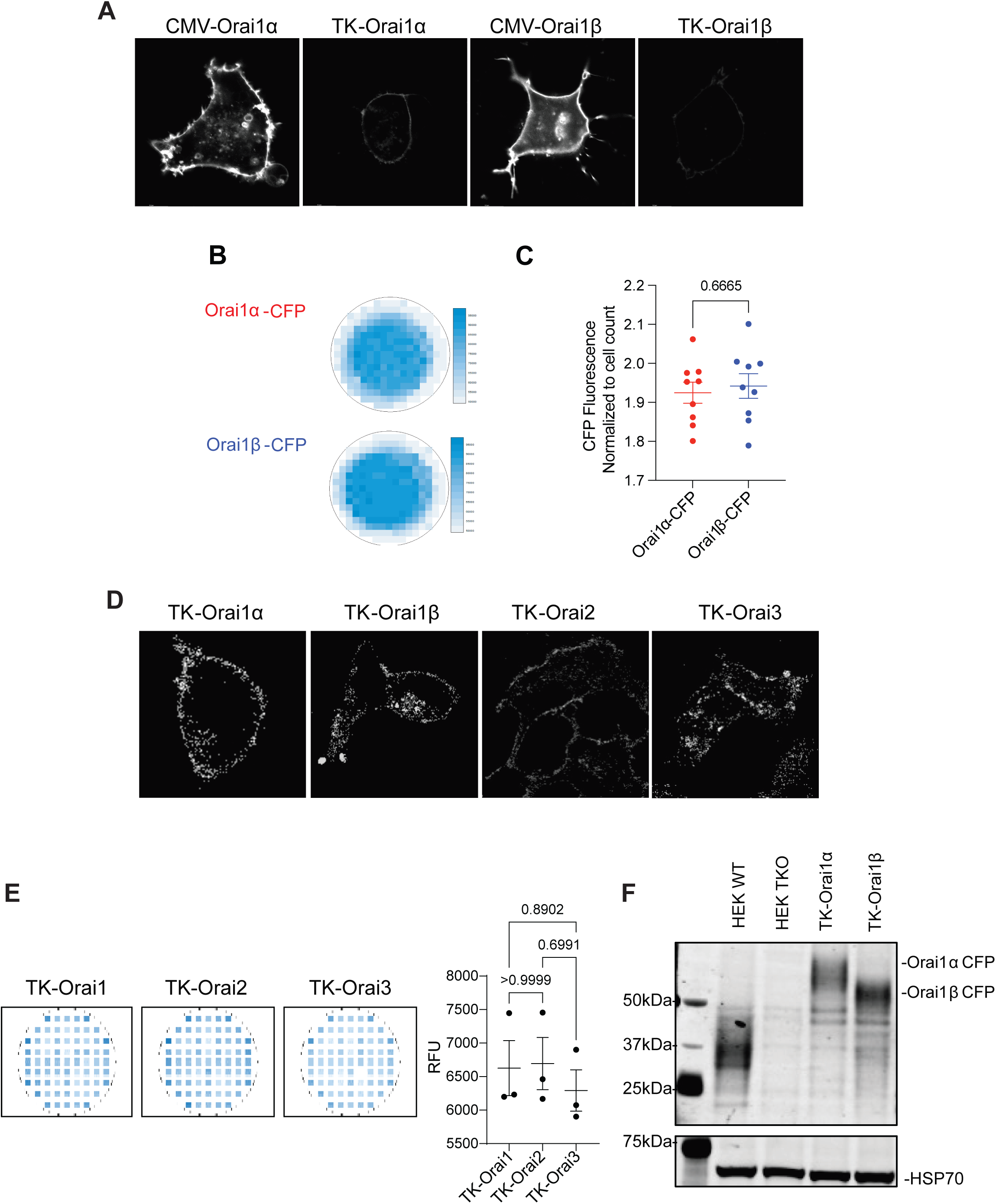
Near-native expression of Orai proteins in Orai-TKO HEK293 cells. **(A)** Representative fluorescence images of HEK293 Orai-TKO cells expressing either Orai1α or Orai1β using TK promoter-driven expression plasmids. Also shown are images of HEK293 Orai-TKO cells expressing Orai1α and Orai1β from the stronger cytomegalovirus (CMV) promoter. **(B-C)** Comparison of CFP fluorescence intensity in cells transfected with Orai1α-CFP or Orai1β-CFP driven by the TK promotor, showing comparable expression between isoforms. **(D)** Confocal fluorescence images of Orai-TKO cells transfected with Orai1α, Orai1β, Orai2, and Orai3 using TK promoter-driven expression plasmids confirming low expression levels. **(E)** Comparison of YFP fluorescence intensity in Orai-TKO cells transfected with Orai1-YFP, Orai2-YFP, or Orai3-YFP driven by the TK promotor. **(F)** Western blot showing endogenous Orai1 in HEK293 cells, lack of Orai1 expression in HEK293 Orai-TKO cells, and ectopic expression of Orai1α-CFP or Orai1β-CFP in HEK293 TKO cells after transfection. Orai1α-CFP and Orai1β-CFP migrate at higher molecular weights than endogenous Orai1 due to CFP tagging, with Orai1α-CFP appearing above Orai1β-CFP, consistent with its longer N-terminus. HSP70 was used as a loading control.

**Figure S2.**
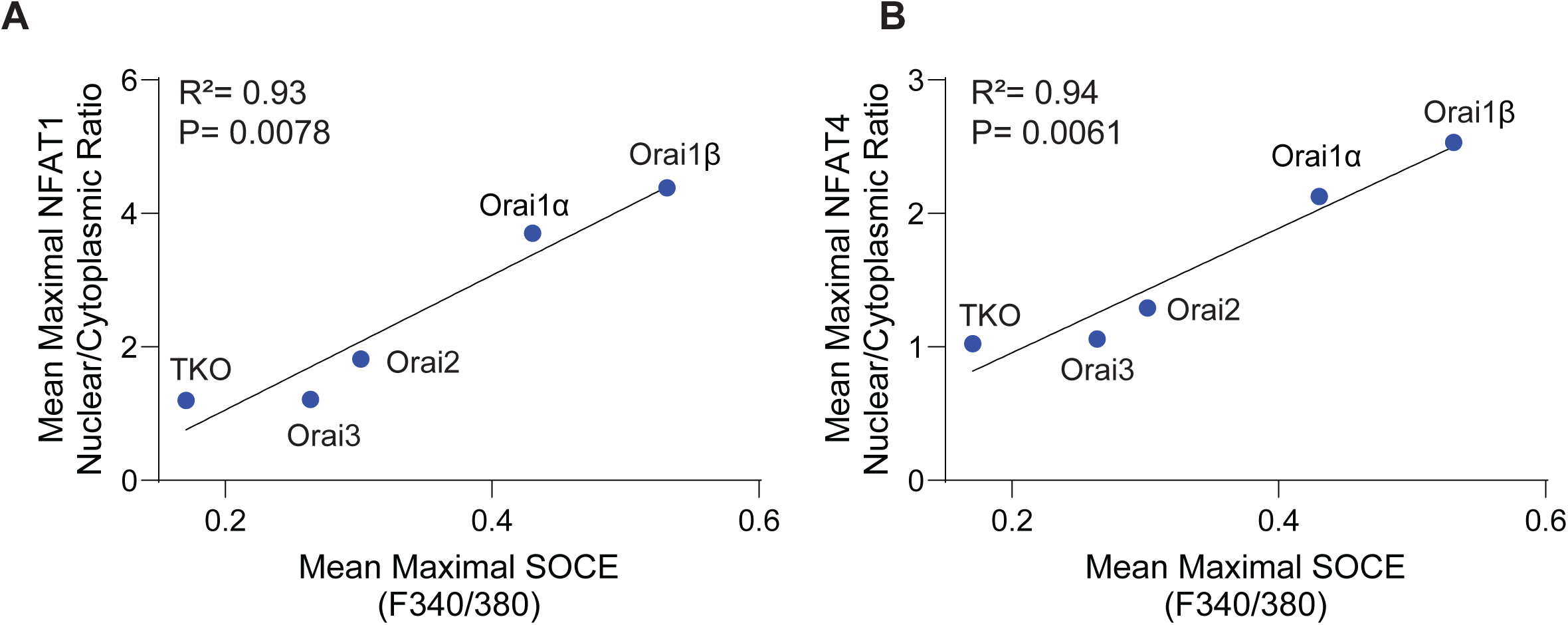
Correlation of SOCE magnitude and NFAT nuclear translocation. Orai-TKO HEK293 cells were left untransfected (TKO) or transfected with either Orai1α, Orai1β, Orai2, or Orai3 using TK promoter-driven expression plasmids. Mean maximal SOCE was plotted against mean maximal **(A)** NFAT1 or **(B)** NFAT4 nuclear translocation for all five experimental conditions. The x-axis value represents the mean maximal Fura-2 ratio (F340/380) measured for SOCE, and the y-axis value represents the corresponding mean maximal NFAT nuclear-to-cytoplasmic ratio. Linear regression showed a strong positive correlation for NFAT1 (R² = 0.93) and NFAT4 (R² = 0.94). SOCE and NFAT measurements were obtained from separate, unpaired experiments.

**Figure S3.**
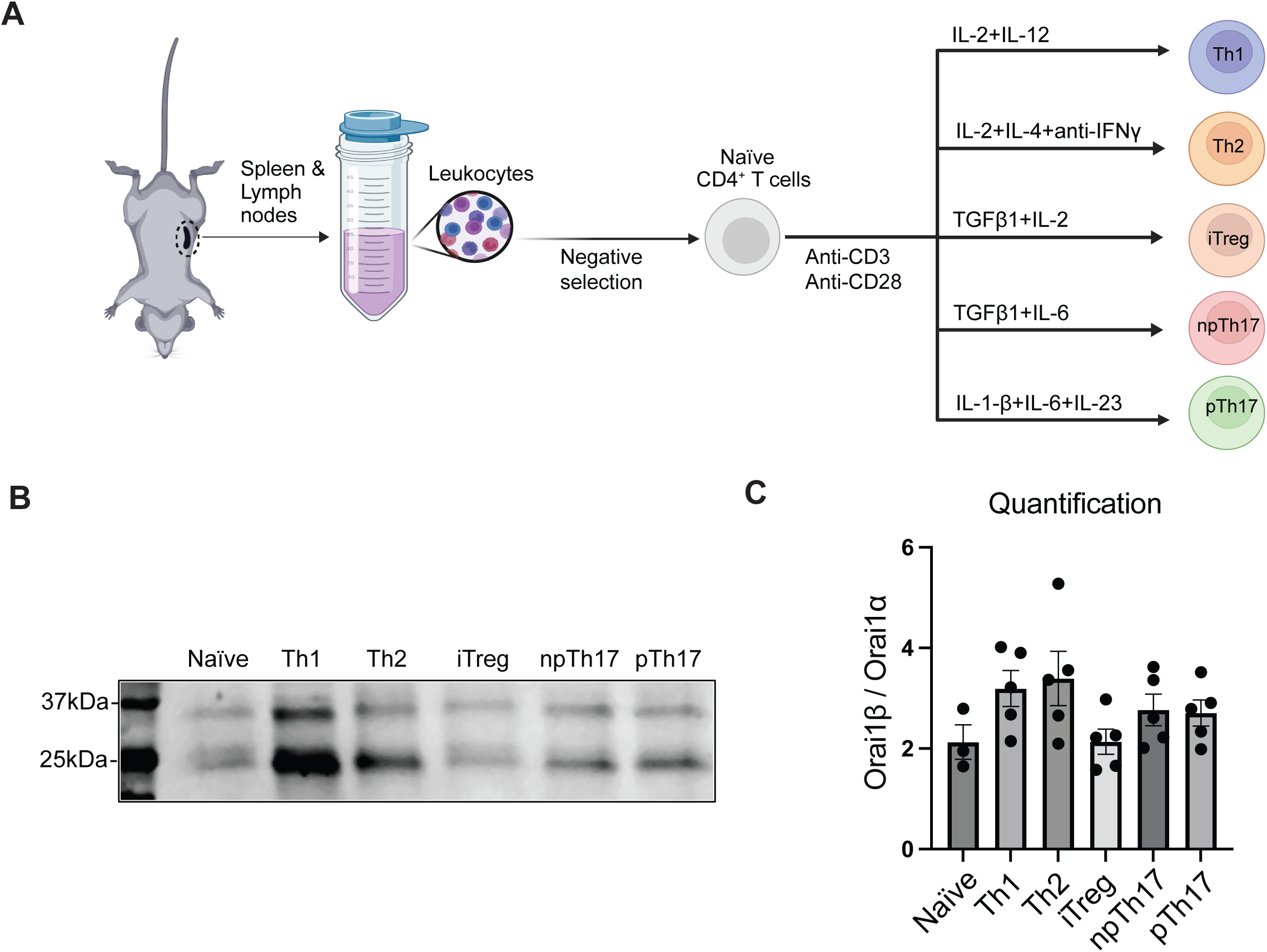
Endogenous expression levels of Orai1α and Orai1β in polarized CD4⁺ T-cell subsets. **(A)** Naïve CD4⁺ T cells were isolated from mouse spleen and lymph nodes by negative selection and activated with anti-CD3/CD28 under lineage-skewing conditions to generate T helper 1 (Th1), Th2, induced Treg (iTreg), non-pathogenic (np) Th17, and pathogenic (p) Th17 cells. **(B)** Orai1α and Orai1β protein expression was assessed by Western blotting in naïve and polarized CD4⁺ T-cell subsets. **(C)** Quantification of the relative Orai1β/Orai1α expression ratios in different CD4^+^ T cell subsets.

**Figure S4.**
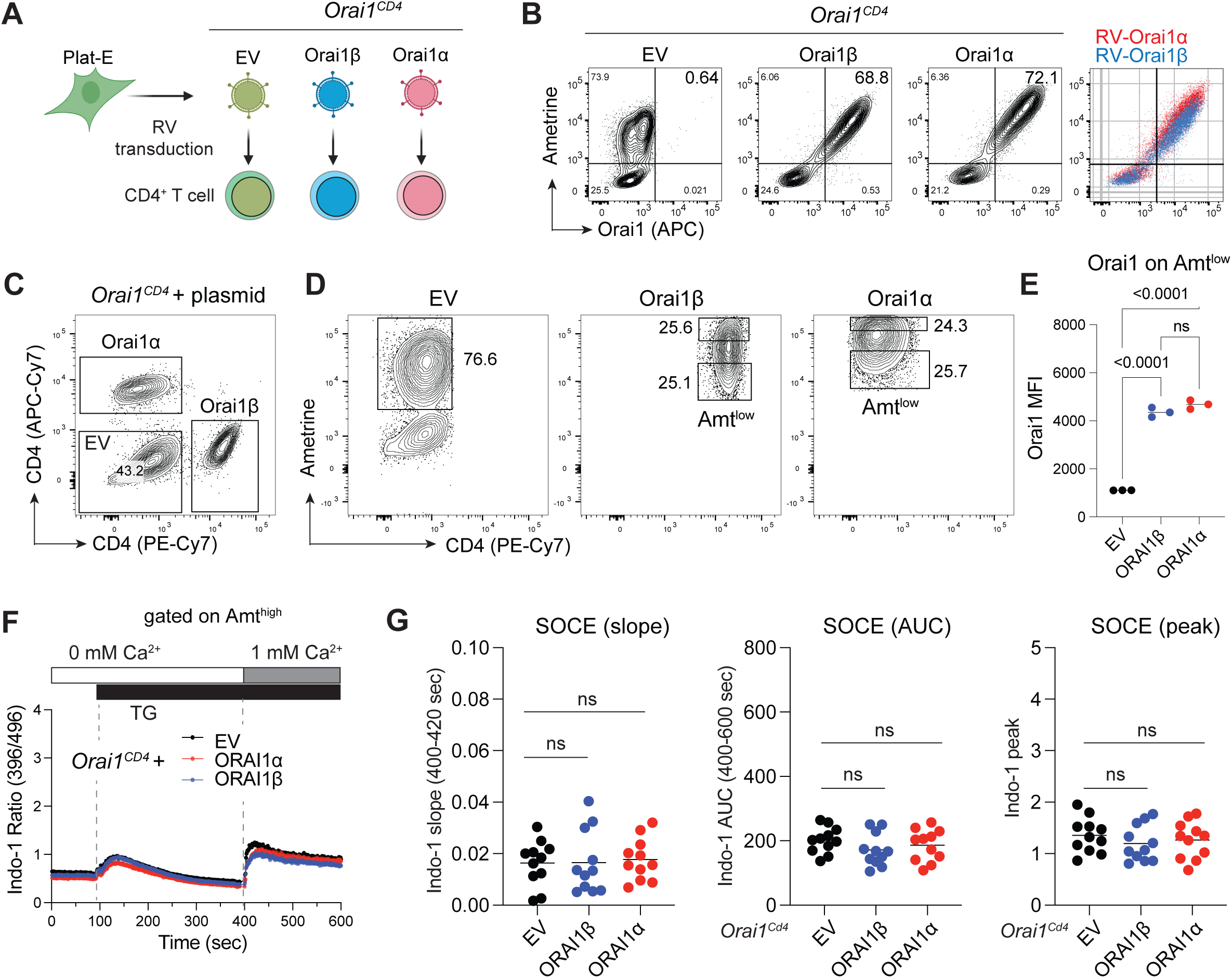
High levels of Orai1α or Orai1β expression suppresses SOCE. **(A)** Retroviral transduction of CD4⁺ T cells from *Orai1^fl/fl^Cd4^Cre^*mice with bicistronic vectors encoding either Orai1α or Orai1β followed by an IRES-Ametrine reporter, or with an empty vector (EV) control. **(B)** Representative flow cytometry plots showing ectopic Orai1 expression (APC, allophycocyanin) in transduced (Amt⁺) CD4⁺ T cells. Fixed and permeabilized cells were stained with a primary anti-Orai1 antibody followed by an Alexa Fluor 647-conjugated secondary antibody. **(C-D)** CD4⁺ T cells from *Orai1^fl/fl^Cd4^Cre^* mice transduced with either Orai1α, Orai1β, or EV and stained with anti-CD4 (APC-Cy7), anti-CD4 (PE-Cy7) or no antibody, respectively, and analyzed for CD4 **(C)** and Ametrine **(D)** expression by flow cytometry. Orai1α and Orai1β expressing CD4⁺ T cells were gated on Amt^high^ (top 25% Ametrine^+^ cells) and Amt^low^ (bottom 25% Ametrine^+^) populations. **(E)** Mean fluorescence intensity (MFI) of Orai1 expression in Amt^low^T cells (from panel D) transduced with Orai1α, Orai1β, or EV. Fixed and permeabilized cells were stained with an anti-Orai1 antibody. **(F)** SOCE measured in Amt^high^ CD4^+^ T cells transduced with Orai1α, Orai1β, or EV and loaded with Indo-1. Cells were stimulated with thapsigargin (TG) in Ca^2+^ free buffer followed by addition of Ca^2+^ containing buffer (final concentration 1 mM Ca^2+^) and analyzed by flow cytometry. **(G)** Quantification of the slope, area under the curve (AUC) and peak of SOCE. Corresponding data for Amt^low^ CD4^+^ T cells are shown in Figure 5A. Data are from 11 mice and four repeat experiments with one technical replicate per condition. Statistical analysis in panels (**E, G**) by ordinary one-way ANOVA multiple comparison. All results are expressed as means ± SEM. ns, not significant (*P* > 0.05).

**Figure S5.**
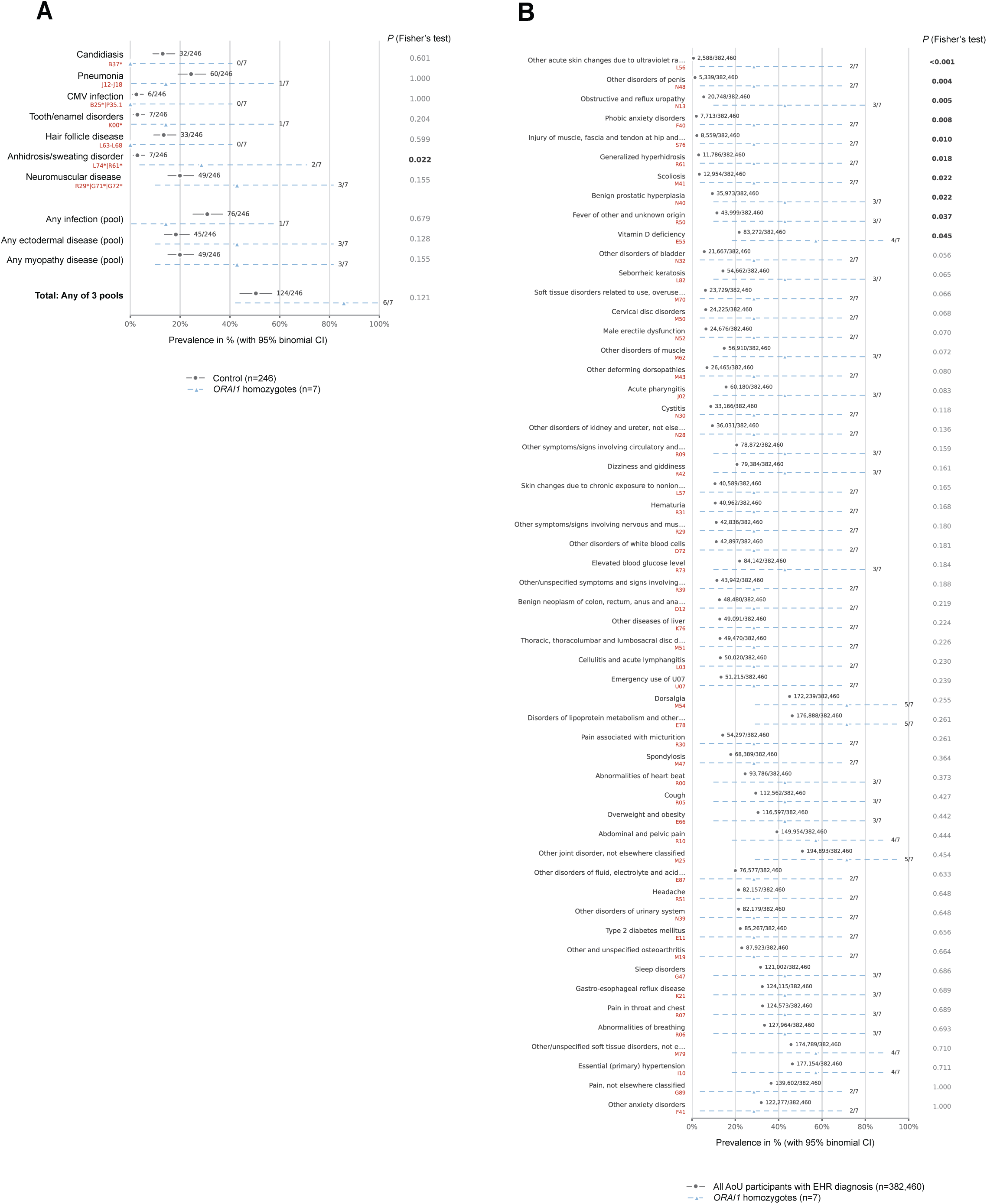
Clinical phenotype analysis of All of Us (AoU) participants homozygous for the Orai1α-specific p.P43TfsX45 variant. **(A)** Prevalence of CRAC channelopathy-associated diagnostic categories in the 7 individuals homozygous for the Orai1α p.P43TfsX45 variant compared to the control group (246 individuals). **(B)** Unbiased exploratory prevalence analysis of broader diagnostic codes in the 7 homozygous individuals compared to the entire AoU cohort. Statistical analysis of significant enrichment of disease codes in A and B was conducted by Fisher’s test.

**Figure S6.**
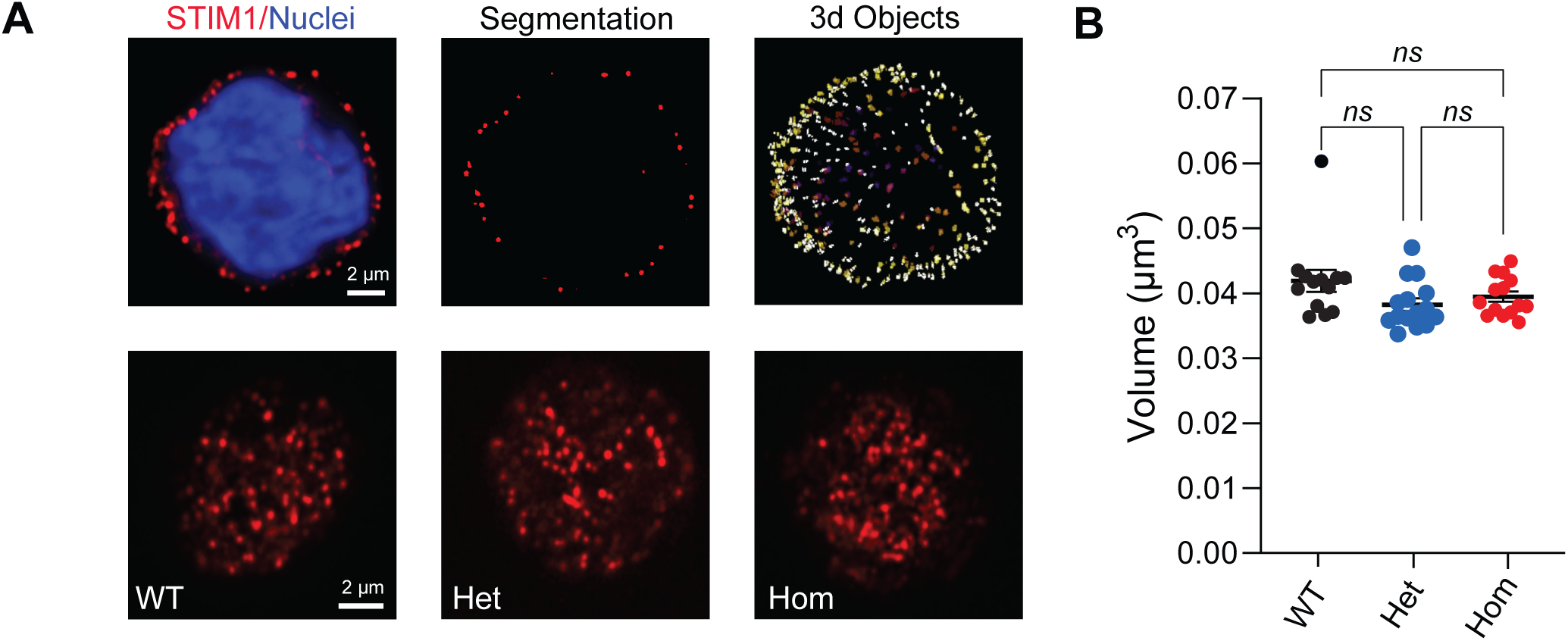
STIM1 clusters in T cells from wildtype and Orai1α null heterozygote and homozygote individuals. **(A)** STIM1 clusters in T cells derived from individuals heterozygous (het) or homozygous (hom) for the Orai1α p.P43TfsX45 variant, or wild-type (WT) controls. Top row: Airy Scan images of STIM1 clusters induced by thapsigargin stimulation analyzed using 3D Spot Segmentation and the volume of each cluster measured. Bottom row: typical examples of the cluster pattern at the bottom of the cells recorded in the three genotypes. **(B)** Summary of the average cluster volume plotted on a per cell basis in the three genotypes. Data are shown as Mean ± SEM with statistical significance assessed using one-way ANOVA followed by Dunnett post-hoc test.

**Figure S7.**
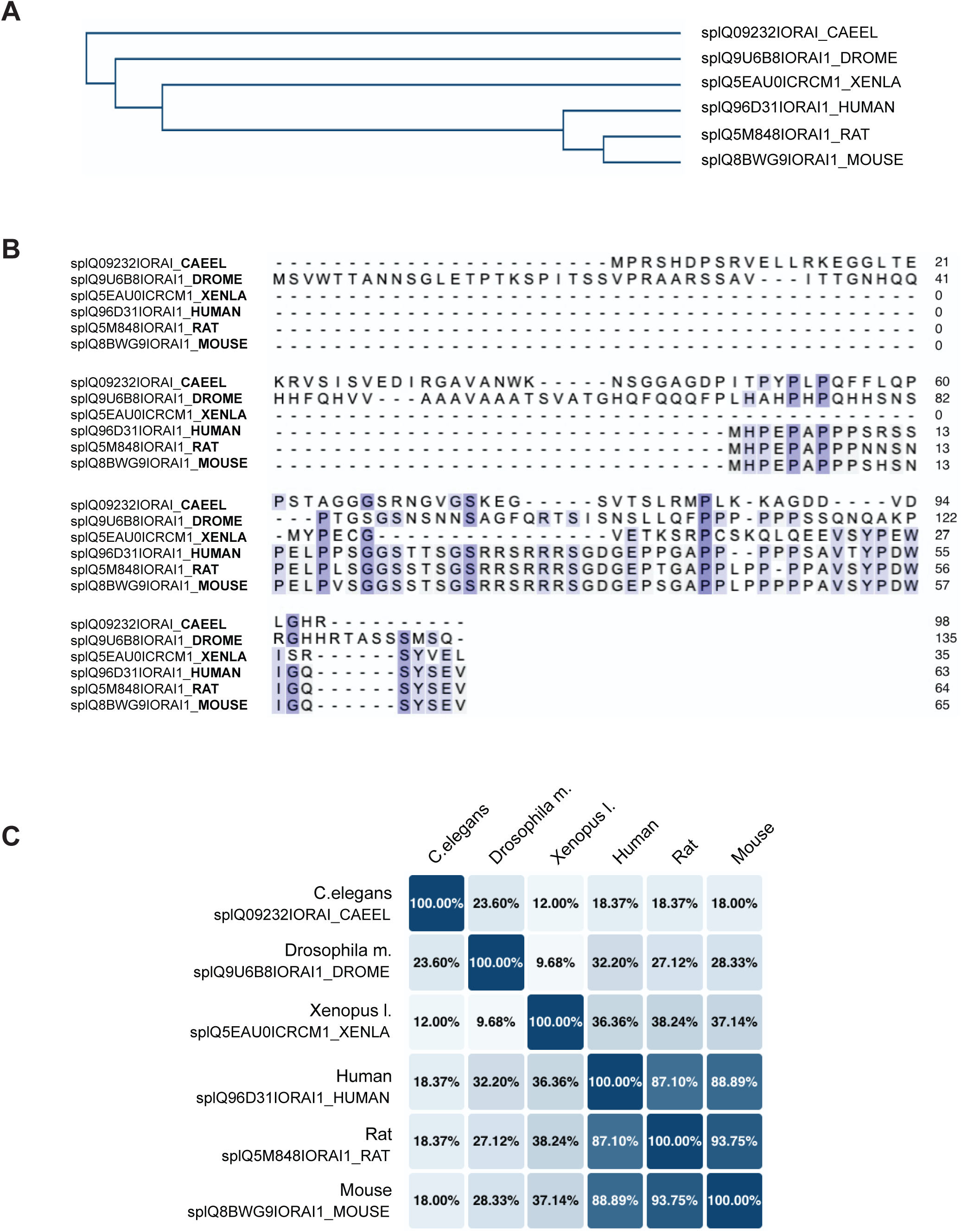
The Orai1α-specific N-terminus has low evolutionary conservation. **(A, B)** Phylogenetic tree **(A)** and multiple sequence alignment **(B)** showing the sequence similarity of the Orai1α-specific N-terminus (amino acids 1–63 of human Orai1) across different species. Sequences were aligned using UniProt alignment tools. Conserved residues are highlighted. **(C)** Percent identity matrix showing pairwise sequence identities for the Orai1α-specific N-terminus across different species.

## Supplementary Table

**Table S1.**
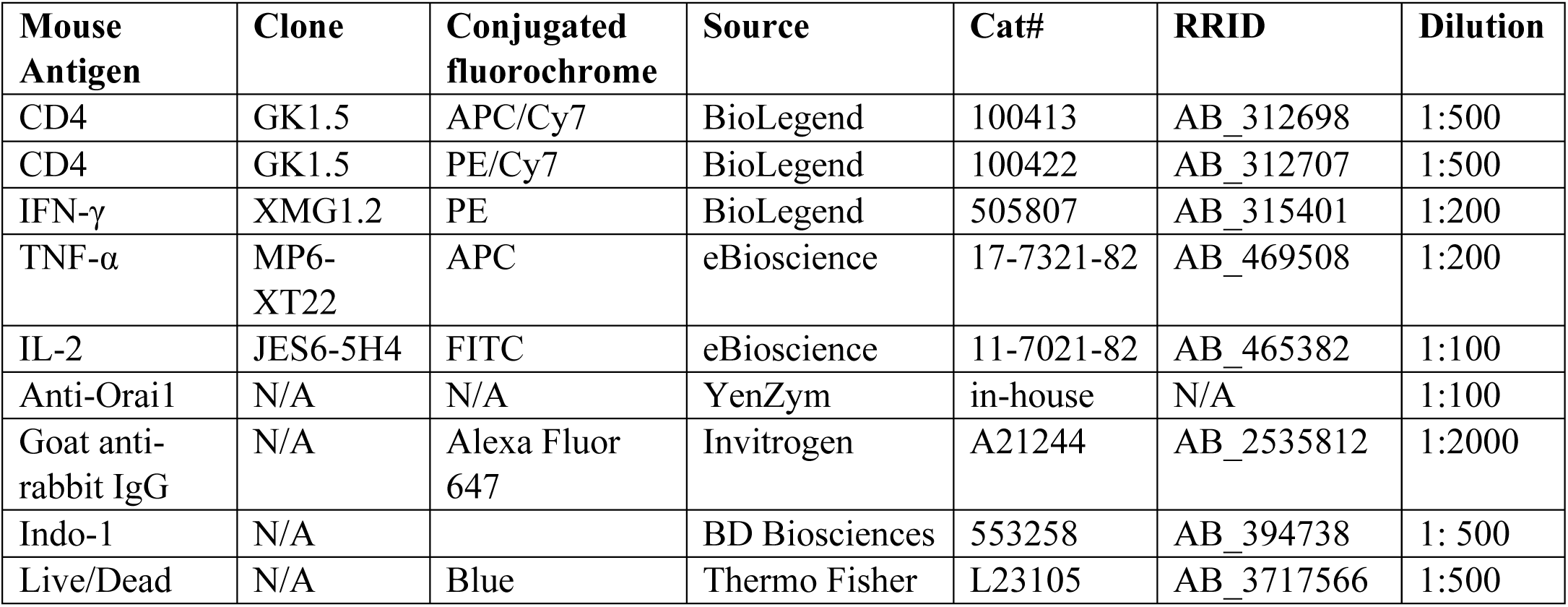
Antibodies used for flow cytometric analysis of murine T cells.

